# BonZeb: Open-source, modular software tools for high-resolution zebrafish tracking and analysis

**DOI:** 10.1101/2021.03.02.433632

**Authors:** Nicholas C. Guilbeault, Jordan Guerguiev, Michael Martin, Isabelle Tate, Tod R. Thiele

## Abstract

We present BonZeb – a suite of modular Bonsai packages which allow high-resolution zebrafish tracking with dynamic visual feedback. Bonsai is an increasingly popular software platform that is accelerating the standardization of experimental protocols within the neurosciences due to its speed, flexibility, and minimal programming overhead. BonZeb can be implemented into novel and existing Bonsai workflows for online behavioral tracking and offline tracking with batch processing. We demonstrate that BonZeb can run a variety of experimental configurations used for gaining insights into the neural mechanisms of zebrafish behavior. BonZeb supports head-fixed closed-loop and free-swimming virtual open-loop assays as well as multi-animal tracking, optogenetic stimulation, and calcium imaging during behavior. The combined performance, ease of use and versatility of BonZeb opens new experimental avenues for researchers seeking high-resolution behavioral tracking of larval zebrafish.

## Introduction

The ability to precisely track animal movements is vital to the goal of relating the activity of the nervous system to behavior. The combination of precision tracking with systems for behavioral feedback stimulus delivery can provide valuable insights into the relationship between sensory inputs and behavioral outputs^1–3^. Methods for high-resolution behavioral tracking and dynamic sensory feedback can be combined with methods for monitoring or manipulating neural circuit activity to allow researchers to explore the neural circuitry underlying sensorimotor behaviors. These assays can also increase the throughput of behavioral experiments through the rapid and repeatable delivery of stimuli, and allow researchers to probe the sensorimotor loop by controlling and perturbing sensory feedback^4–6^.

Despite its significance, the use of behavioral feedback technology is not standard across laboratories investigating sensorimotor integration. A key challenge in developing behavioral feedback systems is synchronizing high-throughput devices. Relatively few software implementations have succeeded in addressing this synchronization problem and most rely on software configurations that require advanced levels of technical expertise. Bonsai, an open-source visual programming language, has recently gained traction among the neuroscience community as a powerful programming language designed for the rapid acquisition and processing of multiple data streams^7^. Currently, there are no released Bonsai packages that allow for high-speed kinematic tracking of small model organisms, such as the larval zebrafish, while providing behavior based stimulus feedback. Here, we present BonZeb, an open-source, modular software package developed entirely in Bonsai for high-resolution zebrafish behavioral tracking with virtual open-loop and closed-loop visual stimulation.

## Results

### Overview

BonZeb provides an open-source and approachable method to implement the functionalities of extant behavioral feedback systems. Furthermore, BonZeb supplies a range of novel utilities that extend Bonsai’s capabilities (**Figure 1**). We developed packages for high-speed video acquisition, presentation of a visual stimulus library, high-resolution behavioral tracking, and analysis. BonZeb inherits Bonsai’s reactive programming framework for processing synchronous and asynchronous data streams (**Figure 2A**). The reactive framework allows users to process incoming streams of data, regardless of whether the data are coming from a finite source (*e.g.* a saved video) or continuous source (*e.g.* streaming from a camera). There are four major classifications of nodes – source nodes, transform nodes, sink nodes, and combinators – which generate or manipulate streams of data called observable sequences (**Figure 2B**). **Figure 2B** also provides a basic example of how BonZeb performs online behavioral tracking. An online manual for BonZeb provides details for running the presented packages as well as a knowledge base for the development of new complex tracking assays (https://github.com/ncguilbeault/BonZeb).

**Figure 1.**
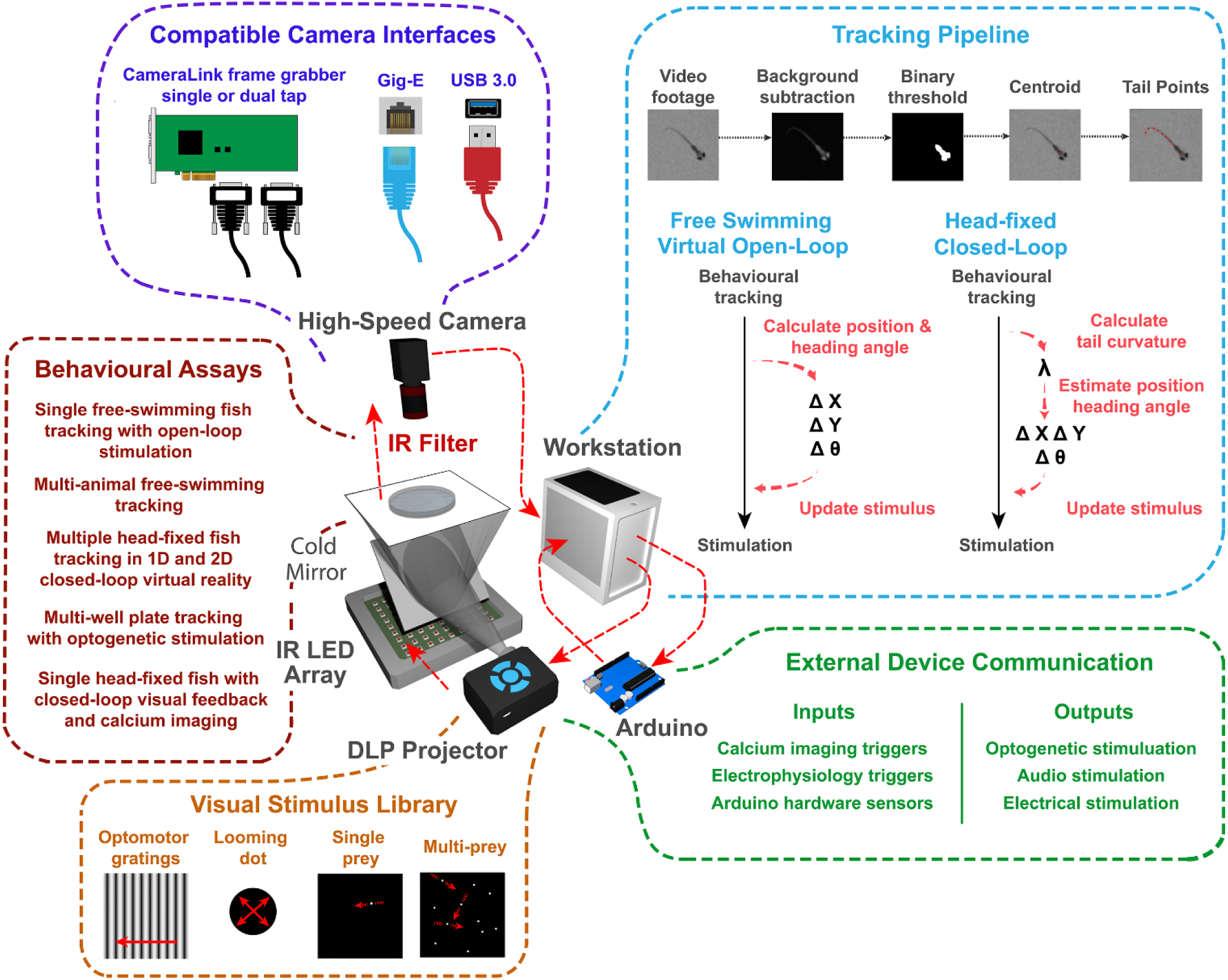
Overview of the behavioral feedback system. The behavioral feedback hardware consists of modular components based on a previous design^5^. High-speed cameras convey live video of behaving larval zebrafish to a workstation, which tracks the behavior of the fish and generates output signals to an external device to provide sensory stimulation. The workstation processes incoming video frames with BonZeb’s customizable tracking pipeline. This pipeline transforms behavioral data into virtual open-loop stimulation for free-swimming fish or closed-loop stimulation for head-fixed fish. BonZeb can interface with Arduino boards, display devices, data acquisition boards, etc., for receiving or sending data. BonZeb also includes a library of common visual stimuli for closed-loop and open-loop visual stimulation. Components of the behavioral feedback system are not to scale.

**Figure 2.**
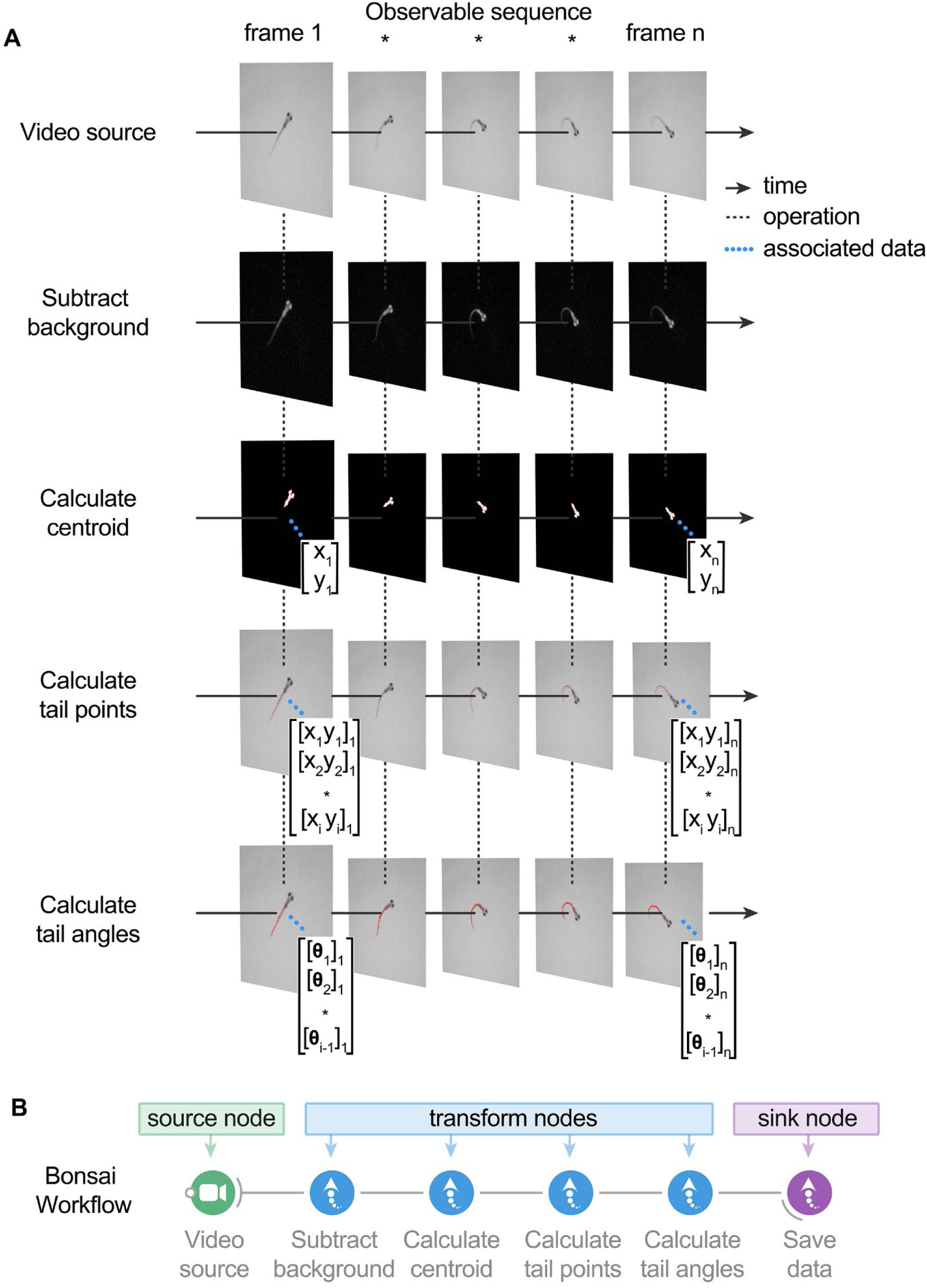
BonZeb inherits Bonsai’s reactive architecture for processing data streams. **(A)** A video source node generates images over time. The video source can either be a continuous stream of images from an online camera device or a previously acquired video with a fixed number of frames. A series of transformation nodes are then applied to the original source sequence. Each transformation node performs an operation on the upstream observable sequence to produce a new observable sequence to downstream nodes. A typical pipeline consists of background subtraction, centroid calculation, tail point calculation, and finally, tail angle calculation. Nodes have a unique set of visualizers that provide the node’s output at each step. Each node has a set of properties associated with the output, such as a single coordinate, an array of coordinates, or an array of angles, which can be used for more sophisticated pipelines. **(B)** Bonsai workflow implementation of the above data processing pipeline. An additional node is attached at the end of the workflow to save the tail angles data to a csv file on disk. There are 4 different general classifications of nodes in Bonsai. Source nodes (green) generate new observable sequences and do not require inputs from upstream nodes. Transform nodes (blue) perform an operation on the elements of an upstream observable sequence to produce a new observable sequence that can be subscribed to by downstream nodes. Sink nodes (purple) perform side operations with the elements of the data stream, such as saving data to disk or triggering an external device. Sink nodes then pass along the upstream observable sequence to subscribed downstream nodes without modifying any of the elements of the upstream observable sequence. Combinator nodes (not shown here) are important for combining sequences and become crucial for more complex pipelines (see online manual for examples).

### Free-Swimming Behavior in Virtual Open-Loop

BonZeb can visually evoke and capture the core behavioral repertoire of freely-swimming larval zebrafish using virtual open-loop assays^4^. To evoke predator avoidance visual escapes, larval zebrafish were presented from below with an exponentially expanding looming dot stimulus to either the left or right visual field at a 90° angle relative to the heading direction (**Figure 3A**). Similar to a previous study^8^, we found that fish stimulated with a looming dot in virtual open-loop produced escape responses characterized by a large initial turn away from the stimulus followed by high-frequency tail oscillations (**Figure 3A, Figure 4A**, Supplemental Video 1: https://git.io/JJABI, Supplemental Video 2 - 700 Hz acquisition: https://git.io/JtMIo). These escape responses were easily identified from other non-escape responses, as the max initial heading angle was consistently greater than 50° and the max velocity exceeded 15 cm/s in all elicited escape responses (**Figure 4A**, bottom left). The max bending angle of the initial turn of the escape depended on the location of the looming stimulus such that the initial turn consistently oriented the fish away from the stimulus (left loom escapes, *n* = 5, *M* = 100.7, *SD* = 27.3, right loom escapes, *n* = 6, *M* = −115.3, *SD* = 33.4, two-tailed t-test: *t*_10_ = 10.47, *p* < 0.001; **Figure 4A**, bottom right).

**Figure 3.**
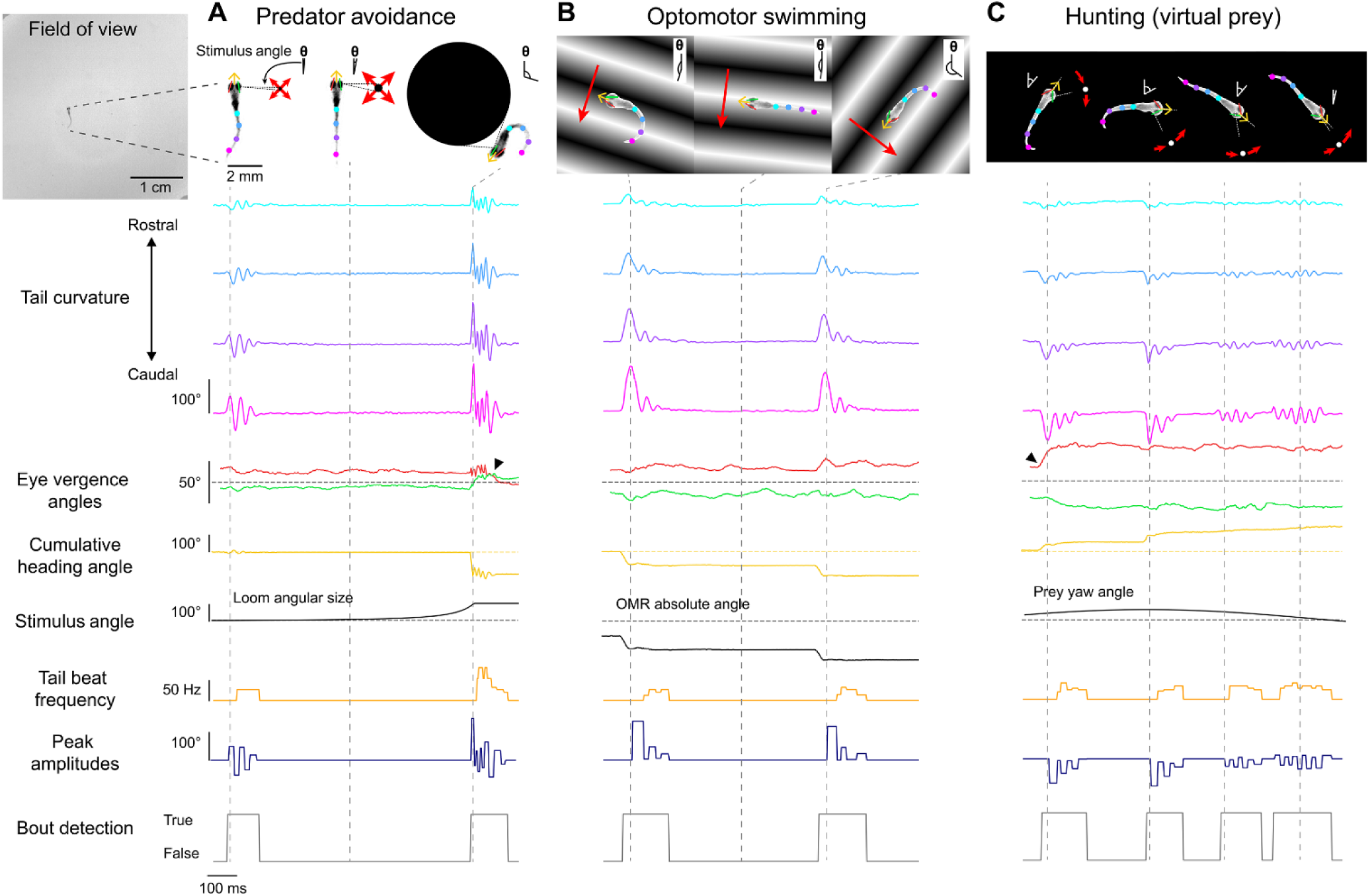
Free-swimming open-loop behavior. Individual freely-swimming zebrafish larva were presented with virtual open-loop visual stimuli while multiple behavioral metrics were recorded (332 Hz). The curvature along 14 tail segments, from the most rostral portion of the tail at the tail base to the most caudal portion of the tail at the tail tip, were calculated and averaged into 4 consecutive bins (cyan to magenta). The angle of the left (red) and right eye (green), the cumulative heading angle (yellow), the visual stimulus angle (black), tail beat frequency (orange), peak amplitudes (navy blue), and bout detection (gray) are displayed. **(A)** A looming dot stimulus (predator avoidance) produced a rapid escape turn followed by a burst swim and a marked divergence of the eyes (arrow). The location of the looming stimulus was fixed with respect to the heading direction and centroid position of the larvae such that the visual angle of the stimulus increased exponentially to a fixed size of the visual field. **(B)** During optomotor swimming, the direction of the OMR stimulus was fixed to the heading direction of the larvae and the point at which the OMR stimulus pivoted was fixed to the larva’s centroid. In this example, the OMR stimulus traversed 90° to the left of the heading direction of the fish, which consistently drove the fish to produce routine turns. **(C)** A small white dot on a black background was presented to larvae from below to create virtual prey stimuli. In this example, the prey stimulus moved along an arc with a fixed radius from the centroid. The velocity of the dot along the arc was defined by a sinusoidal function which reached a maximum of 100°/s directly in front of the larvae and reached a minimum of 0°/s at 60° to the left and to the right of the heading direction. Larvae displayed characteristic hunting behavior towards this stimulus by producing J-Turns when the stimulus was presented to the lateral parts of the visual field and slow, approach swims when the stimulus was presented in the frontal field. These hunting episodes were also characterized by convergence of the eyes throughout the hunting episode (arrow). Images of larvae in **A, B** & **C** were adjusted so they standout against the stimulus background.

**Figure 4.**
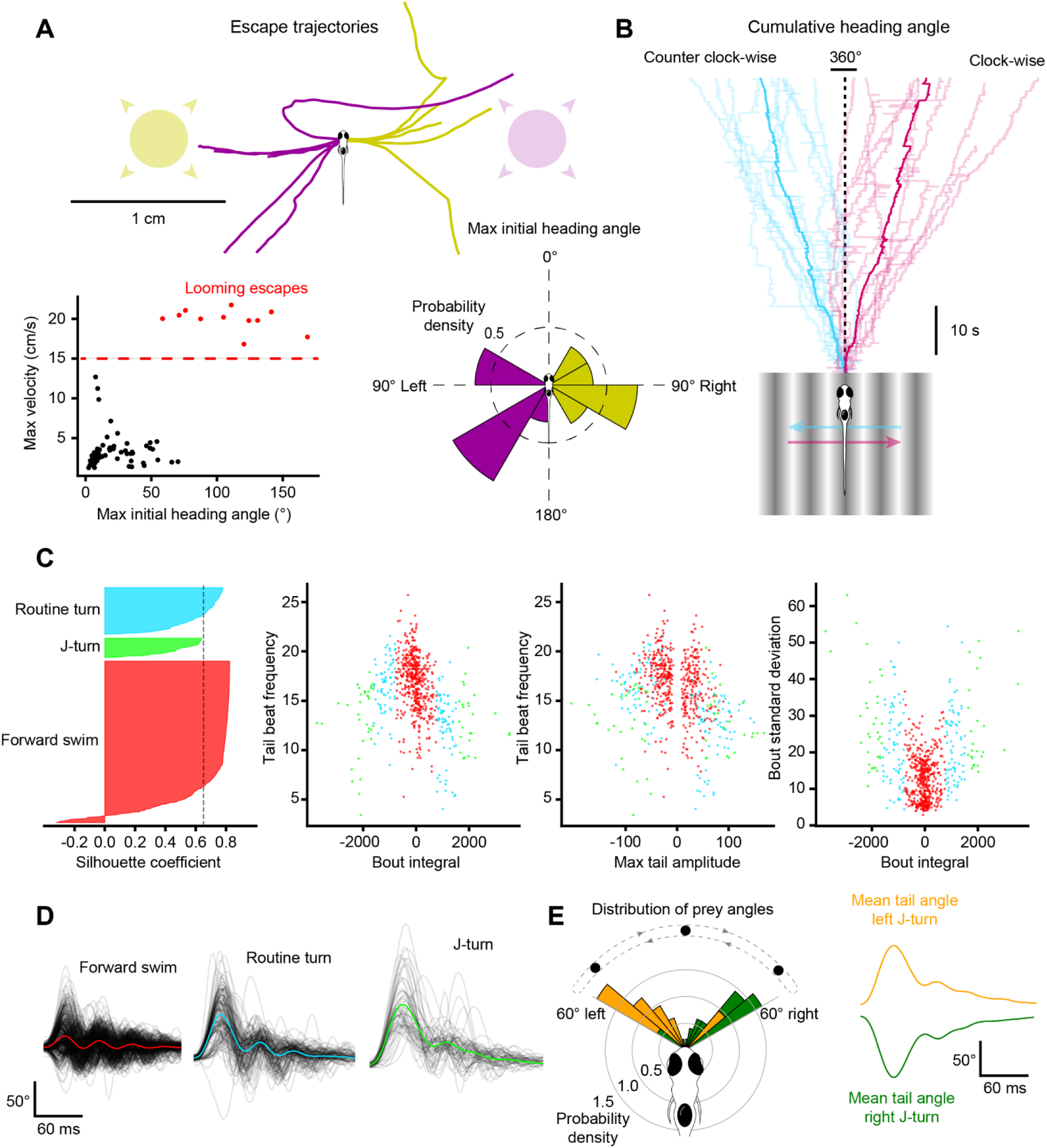
Visual stimulation drives specific behavioral kinematics. **(A)** Escape trajectories in the virtual open-loop looming dot assay when the looming dot was presented from the left (yellow) and from the right (magenta). Bottom left: max velocity (cm/s) with respect to the max initial heading angle (°) plotted for each bout. Bouts classified as escapes are colored red. Dashed red line represents the threshold value applied to the max velocity to select escape responses. Bottom right: probability density distribution for the max initial heading angle. **(B)** Cumulative heading angle over time in response to leftward (blue) and rightward (red) OMR stimulation. **(C)** Hierarchical clustering applied to bouts in response to virtual open-loop prey stimulation. Four kinematics parameters were calculated for each bout (mean tail beat frequency, bout integral, max tail amplitude, bout standard deviation). Left: silhouette plot showing the results of hierarchical clustering with 3 clusters. Dotted line represents silhouette index. Middle-left to right: kinematic parameters plotted for each bout color-coded by cluster. **(D)** Black lines: tail angle over time for every bout in each cluster. Colored lines represent the mean tail angle over time across all bouts in each cluster. Results from **C** and **D** were used to identify the three clusters as forward swims (red), routine turns (blue), and J-turns (green). **(E)** Bouts in the J-turn cluster were separated into left-biased and right-biased swims. Left: probability density distribution of prey yaw angles at the start of left-biased and right-biased swims in the J-turn cluster. Right: mean tail angle over time for left (yellow) and right (green) J-turns.

Fish were stimulated to produce the optomotor response (OMR) under virtual open-loop conditions by continually updating the orientation of a drifting black and white grating. In our open-loop OMR assay, fish were presented with an OMR stimulus from below that was tuned to always drift in a constant direction relative to the heading angle (90° left or right). Consistent with previous studies^5,9^, we observed that fish continually produced turns and swims to follow the direction of optic flow (**Figure 3B**, Supplemental Video 3: https://git.io/JtMIi). Fish produced significantly more clockwise turns when stimulated with rightward OMR (*n* = 10, *M* = 3.8, *SD* = 2.2) compared to when stimulated with leftward OMR (n = 10, *M* = −3.8, *SD* = 1.6, two-tailed t-test: *t*_19_ = 8.42, *p* < 0.001, **Figure 4B**).

We also developed a novel virtual hunting assay where a small white spot is presented from below. **Figure 3C** shows an example of a behavioral sequence from a fish stimulated with a virtual dot moving back and forth along an arc of 120° positioned 5 mm away from the fish. This fish produced two J-turns when the stimulus was in a lateral position followed by two approach swims as the stimulus moved toward the frontal field. In this example, eye tracking was used to detect the characteristic eye convergence that is present in hunting larvae. Supplemental Video 4 (https://git.io/JtyBW) provides an example of a fish engaging in multiple hunting episodes using this stimulus paradigm. We also modified the stimulus such that the stimulus was fixed to one of the lateral extremes of the arc (60^◦^ left or right). When the prey stimulus was fixed to a lateral location in the fish’s visual field, fish often engaged in multiple J-turns with rapid succession. Supplemental Video 5 (https://git.io/JtyBB) provides an example captured at a lower frame rate 168 Hz where a fish performs 6 consecutive J-turns in pursuit of the stimulus.

To determine whether virtual prey stimuli presented in the lateral visual field evoked J-turns, bouts captured during virtual prey stimulation (*n* = 688) were clustered based on four kinematics parameters (bout integral, max bout amplitude, mean tail beat frequency, and bout standard deviation) using hierarchical clustering (**Figure 4C**). A maximum silhouette score was calculated at three clusters (silhouette score = 0.65). Three distinct clusters were detected that resemble the known kinematics of forward swims (*n* = 487), routine turns (*n* = 142), and J-turns (*n* = 59; **Figure 4D**). We examined the location of the prey stimulus when J-turns were produced and observed that the location of the prey stimulus was not uniformly distributed, but highly skewed toward the lateral visual field (**Figure 4E**). Further decomposition of the J-turn cluster into left vs right J-turns revealed that left J-turns (*n* = 34, **Figure 4E**, top right) were produced when the prey was positioned in the far-left visual field whereas right J-turns (*n* = 25, **Figure 4E**, bottom right) were generated in response to prey stimuli located in the far-right visual field (two-tailed t-test: *t*_58_ = 3.72, *p* < 0.001; **Figure 4E**).

### Multi-Animal Tracking with Visual Stimulation

We created a BonZeb protocol to track multiple fish during OMR stimulation. When the center of mass of the group crossed into the leftmost quarter of the arena, the direction of the OMR stimulus updated to move rightwards and vice versa when the center of mass entered the rightmost quarter of the arena. Robust optomotor swimming back and forth across the arena was observed and real-time tracking resolved detailed tail kinematics across all individuals (*n* = 12) (**Figure 5A**, Supplemental Video 6 provides another example: https://git.io/JtyBz). We also developed a multi-animal hunting assay where larvae (*n* = 6) were tracked while a group of moving virtual prey were projected from below. When presented with virtual prey, larvae performed characteristic J-turns and slow approach swims (**Figure 5B**).

**Figure 5.**
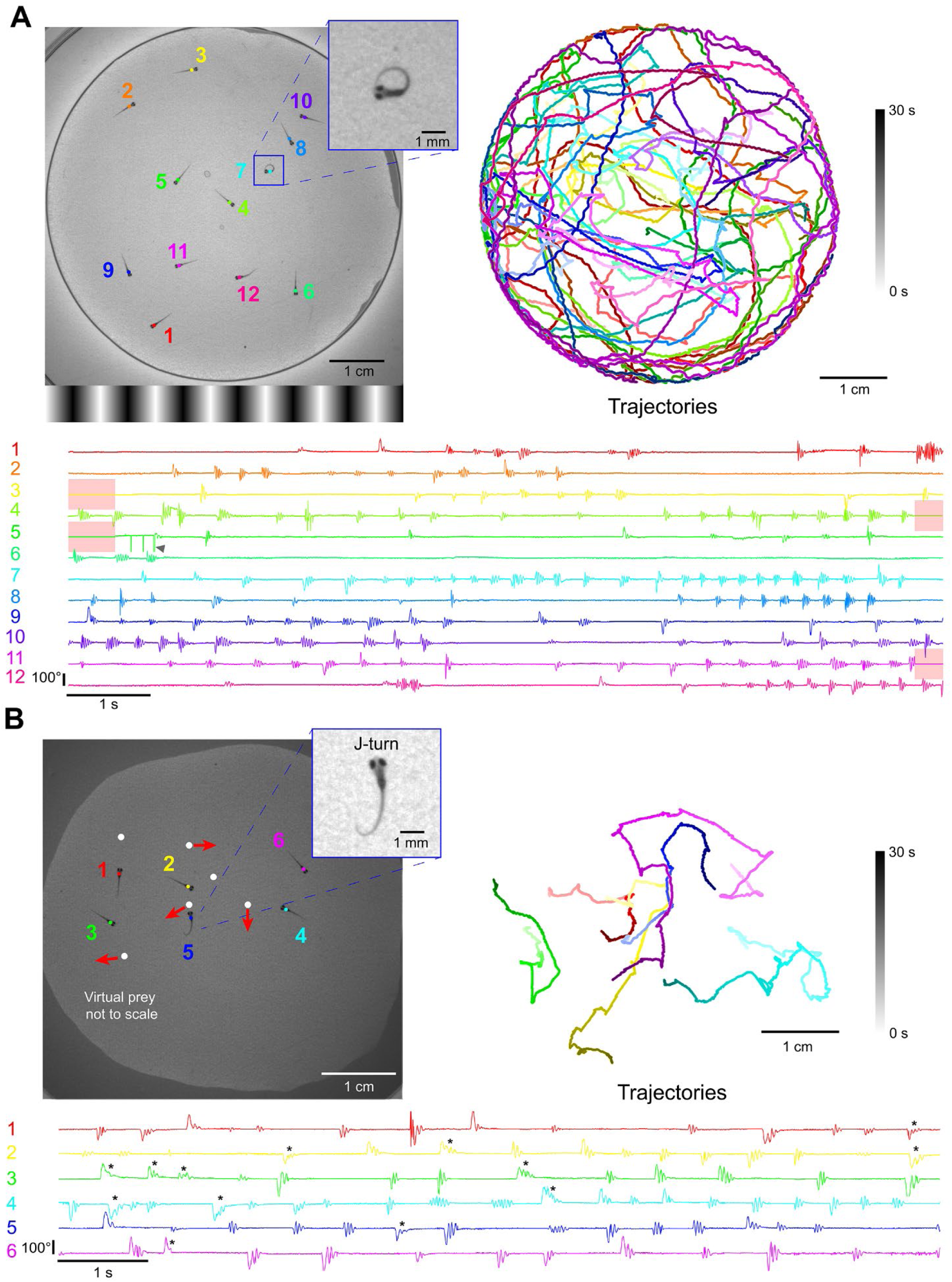
Multi-animal tracking during OMR and prey capture. **(A)** The position and tail curvature of 12 larvae tracked (332 Hz) during the presentation of an OMR stimulus. The direction of the OMR stimulus changed to traverse leftward or rightward depending on whether the center of mass of the group crossed into the rightmost quarter or leftmost quarter of the arena, respectively. Trajectories are individually color coded by fish and shaded by time. The tail curvature for each fish is represented as the average of the 3 most caudal tail segments. Post processing of the tail curvature data revealed sections of the data when fish physically contacted each other (red highlighted regions). The tail tracking results surrounding these encounters decreased in accuracy and occasionally produced tracking errors (arrow). **(B)** Freely swimming larvae in a group of 6 were tracked and presented with multi-prey stimuli projected from below. Virtual prey were programmed to produce paramecia-like behavior, defined as periods of forward movement, brief pauses, and changes in orientation. The linear velocity, distance, length of pause, angular velocity, and orientation were varied for each virtual prey. Larvae produced distinct J-turns in response to virtual prey (* = manually identified J-turn).

We found that fish-to-fish contact, especially at the edges of the arena, led to sporadic swapped identities and tracking errors. To help offset such errors, we automatically discarded regions of the dataset when larvae physically contact each other (**Figure 5A**, red shaded regions). We calculated the percentage of tracking data where no fish-to-fish contacts were detected and found that our method had reliably produced > 90% accuracy with group sizes ranging from 5 – 20 (**Supplemental Figure 1C**). We also sought to determine how the number of fish tracked (centroid and tail kinematics) affected performance. The time between successive tracked frames was normally distributed around the theoretical frame rate (3.01 ms) for a single animal tracked (*M* = 3.03, *SD* = 0.58) and as the number of tracked fish increased, the average time between tracked frames also increased (group size = 5, *M* = 3.27, *SD* = 1.54; group size = 10, *M* = 3.39, *SD* = 2.1; group size = 15, *M* = 3.55, *SD* = 2.25; group size = 20, *M* = 3.49, *SD* = 2.77)(**Supplemental Figure 1B**). Additionally, as the number of fish increased, we observed a greater percentage of values in the range of 9 ms – 23 ms (group size = 1, 0.08%; group size = 5, 0.91%; group size = 10, 1.88%; group size = 15, 2.36%; group size = 20, 3.75%).

### One-Dimensional Closed-Loop Optomotor Swimming

We implemented an experimental protocol that allowed us to present closed-loop optomotor gratings with varying feedback gain constants to head-fixed fish. This experimental design was previously developed to investigate how optomotor swimming adapts to altered visual feedback^10^. With our hardware configuration, the closed-loop round-trip stimulus delay averaged 64.1 ms with an average of 8.3 ms attributed to BonZeb processing and the remainder to stimulus delivery delays imposed by the projector (frame buffering and refresh rate) (**Supplemental Figure 1A**). Consistent with the previous study^10^, fish displayed stark behavioral differences between trials of different gain factors. **Figure 6A** shows representative examples of bouts from the same fish under low (0.5), medium (1.0), and high (1.5) gain conditions. Most notably, bouts were more frequent, longer, and had greater tail beat frequency with low gain conditions compared to medium and high gain conditions (**Figure 6B, Figure 6C**, Supplemental Video 7: https://git.io/JtyB2). We found that fish produced a greater number of swim bouts in low gain, with bout numbers decreasing as the gain increased (*n* = 16, one-way ANOVA: *F*_2,45_ = 20.76, *p* < 0.001; **Figure 6C**, far-left). Bout duration decreased as gain factor increased (*n* = 16, one-way ANOVA: *F*_2,45_ = 16.92, *p* < 0.001; **Figure 6C**, middle left). Fish displayed shorter inter-bout intervals in low gain compared to medium and high gain conditions (*n* = 16, one-way ANOVA: *F*_2,45_ = 29.5, *p* < 0.001; **Figure 6C**, middle right) and mean tail beat frequency decreased with increasing gain values (*n* = 16, one-way ANOVA: *F*_2,45_ = 13.09, *p* < 0.001; **Figure 6C**, far-right).

**Figure 6.**
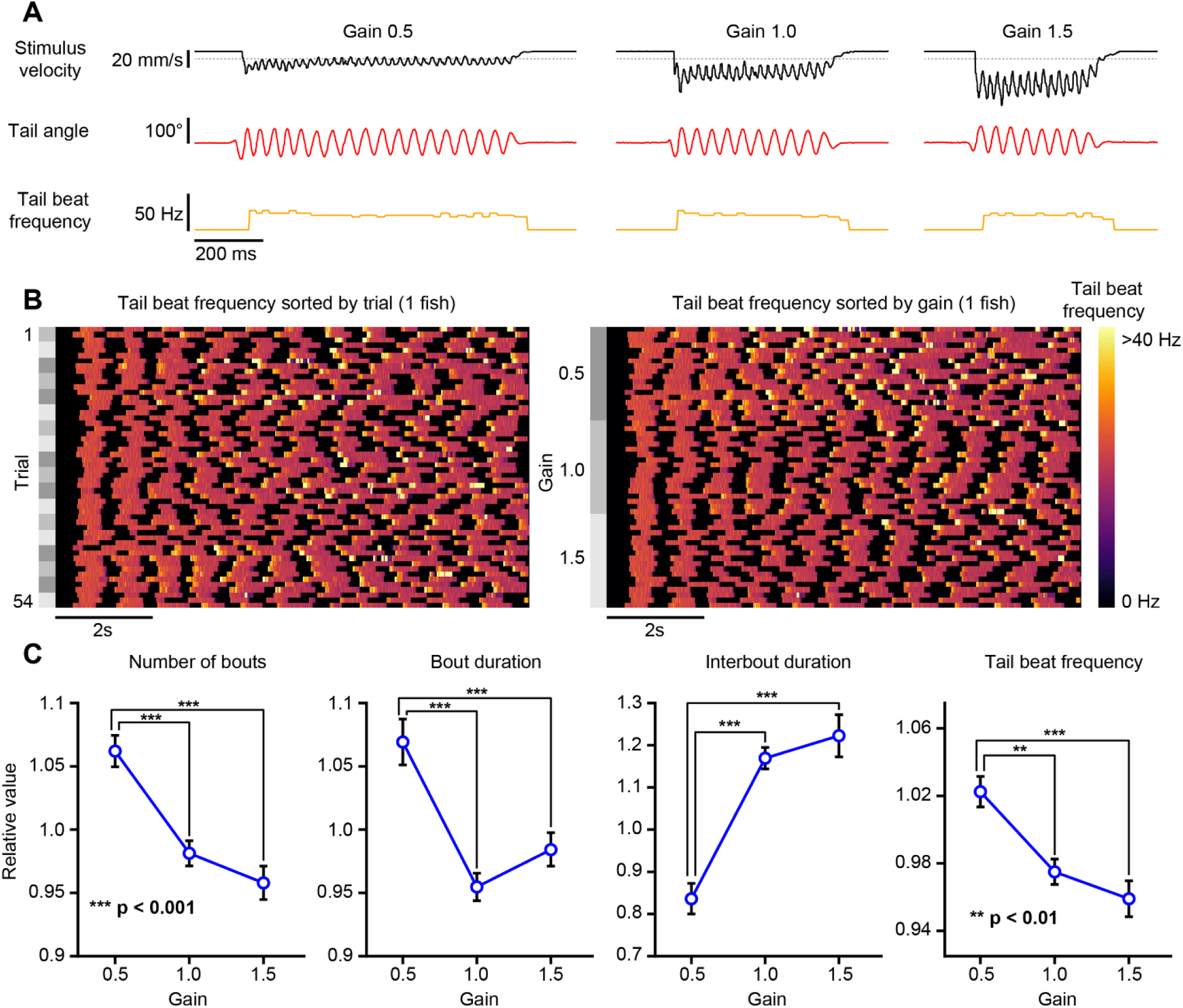
One-dimensional head-fixed closed-loop OMR. The tail of a head-fixed fish was tracked (332 Hz) while presenting a closed-loop OMR stimulus with varying feedback gain values. **(A)** Example bouts taken for each gain value for a single fish. Fish were tested on 3 different gain values. Left: gain 0.5. Middle: gain 1.0. Right: gain 1.5. The stimulus velocity (black), tail angle (red), and tail beat frequency (orange) were recorded continuously throughout the experiment. The stimulus velocity changed in proportion to the swim vigor multiplied by the gain factor. Dotted lines indicate baseline value of 0. **(B)** An example of a single fish showing the tail beat frequency across trials. Left: tail beat frequency sorted by trial number. Right: same data as left with tail beat frequency sorted by gain. **(C)** Bout kinematics plotted as a function of gain factor.

### Multi-Animal Two-Dimensional Closed-Loop Optomotor Swimming

We designed a two-dimensional closed-loop assay which provided both forward-backward, as well as angular visual feedback to multiple head-fixed fish independently. This assay was inspired by a previous implementation of a two-dimensional closed-loop optomotor assay^11^. We found that fish consistently responded to gratings of different orientations by swimming in the direction of optic flow (**Figure 7A**). The orientation of the gratings converged towards the heading angle as trials progressed over 30 seconds (**Figure 7B**). The probability that the final stimulus orientation converged (*P* = 0.62) towards the heading angle was much greater than the probability that the orientation diverged (*P* = 0.28) or remained unchanged (*P* = 0.11). At larger orientations, fish tended to produce turns over forward swims, as indicated by the mean tail angle across swimming bouts. Bouts tended to transition from asymmetric turns in the direction of the stimulus orientation to symmetric forward swims as trials progressed and the orientation of the stimulus neared alignment with the heading angle (*n* = 4668, one-way ANOVA: *F*_6,4661_ = 224.01, *p* < 0.001; **Figure 7C**, left). The mean angular velocity of the stimulus showed a similar trend, albeit in the opposite direction, such that the angular velocity produced by swimming bouts at large stimulus orientations was larger and tended to align the stimulus orientation to the heading direction (*n* = 4668, one-way ANOVA: *F*_6,4661_ = 260.34, *p* < 0.001; **Figure 7C**, right). Tail beat frequency was significantly different for stimulus orientations between −22.5° and 22.5° compared to all other stimulus orientations. Tail beat frequency increased as the orientation of the stimulus became aligned to the heading direction (*n* = 4668, one-way ANOVA: *F*_6,4661_ = 38.7, *p* < 0.001; **Figure 7C**, middle).

**Figure 7.**
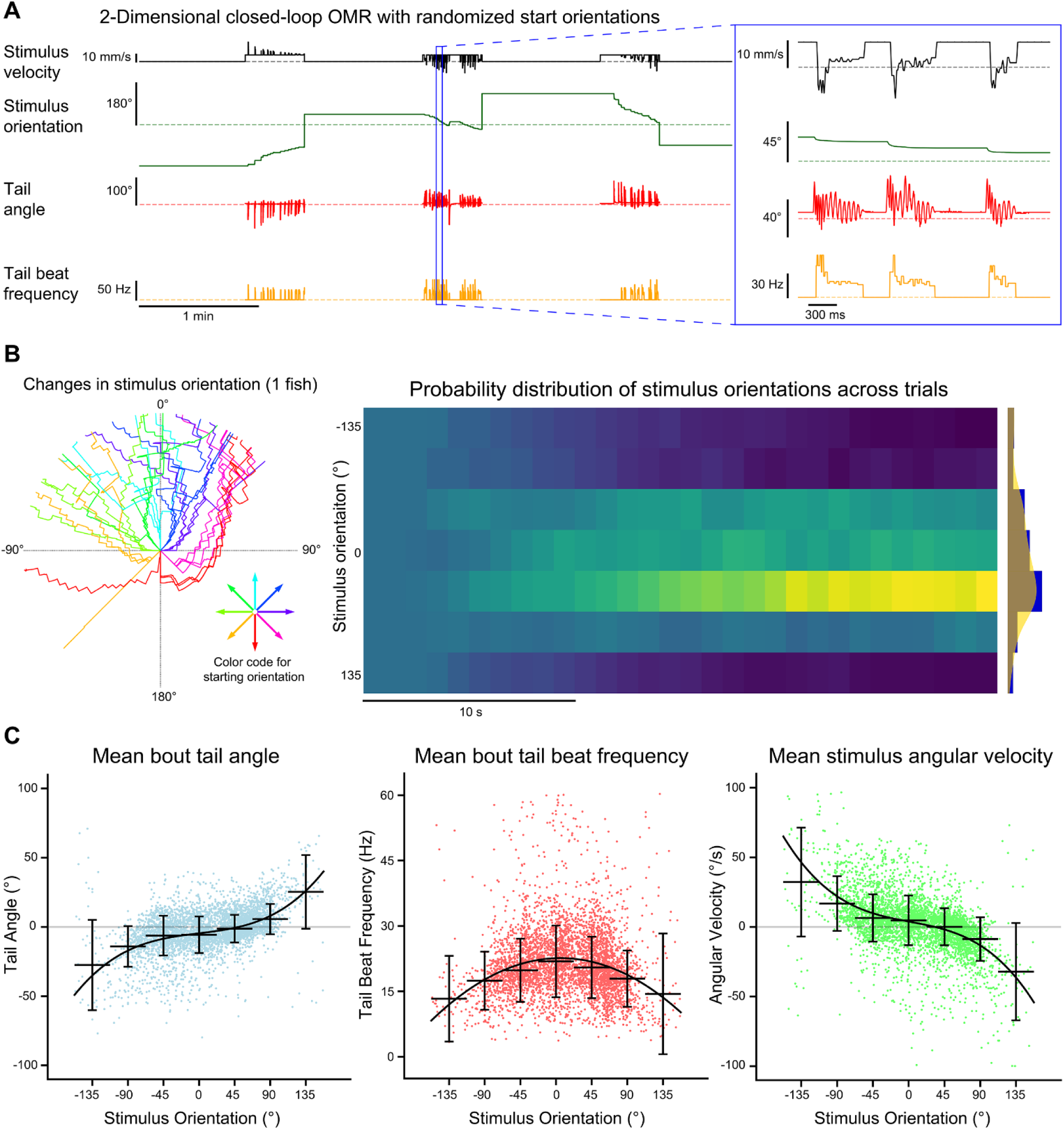
Multi-animal two-dimensional closed-loop OMR. The tails of four head-fixed fish were tracked (332 Hz) while each fish was presented with a two-dimensional closed-loop OMR stimulus. **(A)** Data collected for each fish included the stimulus velocity (black), stimulus orientation (green), tail angle (red) and tail beat frequency (orange). The stimulus velocity changed in proportion to the tail beat frequency and current stimulus orientation. The stimulus orientation changed with the tail angle. Trials were 30 seconds long and were preceded by a 1-minute rest period where only the stimulus velocity and stimulus orientation data were recorded. At the end of each trial, the orientation of the stimulus was reset to one of the 8 randomized start orientations. **(B)** Stimulus orientation over time. Left radial plot: stimulus orientation is plotted for all trials for a single fish. Trials are color-coded based on initial starting orientation and extend outwards from the origin as a function of time. Right heatmap: normalized histograms of stimulus orientations over time across fish (*n* = 16). Binning was 1 second for the x-axis and 45° for the y-axis. Far right plot: histogram and kernel density estimate of the distribution of orientations at the end of trials across all fish. **(C)** Bout kinematics and stimulus angular velocity are plotted for each bout as a function of stimulus orientation at bout onset. Orientations were divided into 45° bins and the median and standard deviation for each bin are plotted. A polynomial function was fit to each dataset. The mean bout tail angle and mean stimulus angular velocity were fit with 3rd degree polynomials (R^2^ = 0.223 and R^2^ = 0.256, respectively) whereas the mean bout tail beat frequency was fit with a 2nd degree polynomial (R^2^ = 0.05).

### Multi-Animal Tracking with Optogenetic Stimulation

To demonstrate how BonZeb can be used for high-throughput behavioral analysis with neural circuit manipulations, we developed a novel multi-animal optogenetic stimulation assay (**Figure 8A**). The setup allowed us to deliver stimulation to multiple fish simultaneously and track each fish’s trajectory over time (**Figure 8B, Figure 8C**). Experimental animals that expressed channelrhodosin-2 in glutamatergic neurons displayed a marked increase in locomotor activity during periods of optogenetic stimulation compared to control animals (**Figure 8D**). We averaged across 2-minute periods within an experiment, starting 30 seconds before and ending 30 seconds after each stimulation period, and found the mean instantaneous velocity of experimental animals to be greater than control animals during stimulation (**Figure 8E**). We examined the total distance travelled when stimulation was ON compared to when stimulation was OFF across the entire experiment for both experimental (*n* = 15) and control animals (*n* = 15). Using a mixed ANOVA, we found a significant effect for group (*F*_1,28_ = 27.42, *p* < 0.001), a significant effect for stimulation (*F*_1,28_ = 33.66, *p* < 0.001), and a significant interaction between group and stimulation (*F*_1,28_ = 45.84, *p* < 0.001). Pairwise comparisons using paired t-tests revealed that for the experimental animals, the total distance travelled for stimulation ON (*M* = 442.85, *SD* = 127.31) was significantly higher compared to stimulation OFF (*M* = 214.73, *SD* = 43.53; *t*_14_ = 6.44, *p* < 0.001), whereas the total distance travelled for the control animals during stimulation ON (*M* = 225.78, *SD* = 32.36) was significantly less than stimulation OFF (*M* = 243.37, *SD* = 21, *t*_14_ = 2.2, *p* = 0.045; **Figure 8F**).

**Figure 8.**
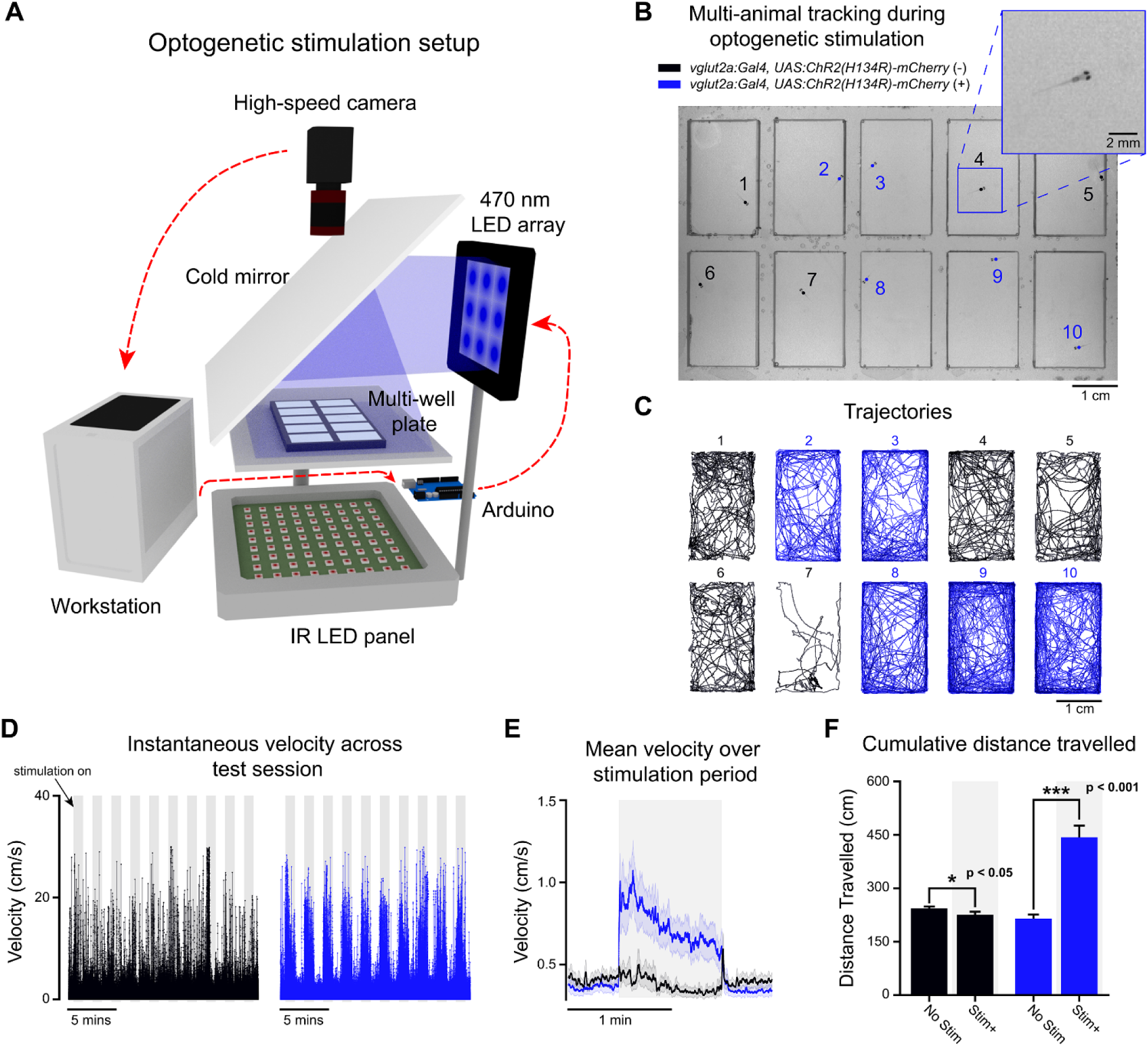
Free-swimming multi-animal tracking during optogenetic stimulation. Ten larvae were tracked simultaneously (200 Hz) in a multi-well plate while being subjected to epochs of optogenetic stimulation. **(A)** 3D schematic of the optogenetic stimulation setup. Live video is processed by BonZeb, which sends commands to an Arduino to generate a digital signal for triggering a 470 nm LED array (0.9 mW/mm^2^ at the plate). Components of the optogenetic setup are not to scale. **(B)** Example of a single video frame. Data are color coded based on genetic background. **(C)** Example of trajectories over an entire 20-minute test session. **(D)** Instantaneous velocities for all fish plotted across the entire test session. Shaded regions represent periods when optogenetic stimulation occurred. 1-minute intervals of stimulation and no stimulation repeated throughout the assay with a 30 second no stimulation period occurring at the beginning and end of the assay. Left: instantaneous velocity of control larvae (*n* = 15). Right: instantaneous velocity of experimental larvae (*n* = 15). **(E)** Mean instantaneous velocity averaged across all fish and all stimulation periods. Instantaneous velocity across each stimulation period was averaged for each fish and convolved with a box car filter of 200 samples. Bold line represents the mean and shaded region represents SEM. (**F**) Cumulative distance travelled during periods of stimulation ON (Stim+) and stimulation OFF (no stim).

### Closed-loop Visual Stimulation during Calcium Imaging

To demonstrate how BonZeb can be used in combination with techniques for imaging neural activity, we implemented a one-dimensional closed-loop OMR assay during two-photon calcium imaging. We performed volumetric calcium imaging in the hindbrain (500 x 500 x 63 µm) of a fish expressing GCaMP6f throughout the brain during OMR stimulation^12^. We imaged at 2.7 Hz using an objective piezo and resonant scanning (9 z-planes, 7 µm spacing). OMR stimulation was presented to one side of the fish using a projector (**Figure 9A**). We found that fish consistently produced swims when forward moving gratings were presented (**Figure 9B**). We used an automated ROI extraction method, based on the CaImAn analysis package^13^, to find 25 neuronal ROIs in the hindbrain for each imaged z-plane recorded (**Figure 9C**). We observed an increase in neuronal activity across planes during swim bouts, in agreement with previous studies where hindbrain activity was shown to be highly correlated with swimming^5,14,15^ (**Figure 9D**).

**Figure 9.**
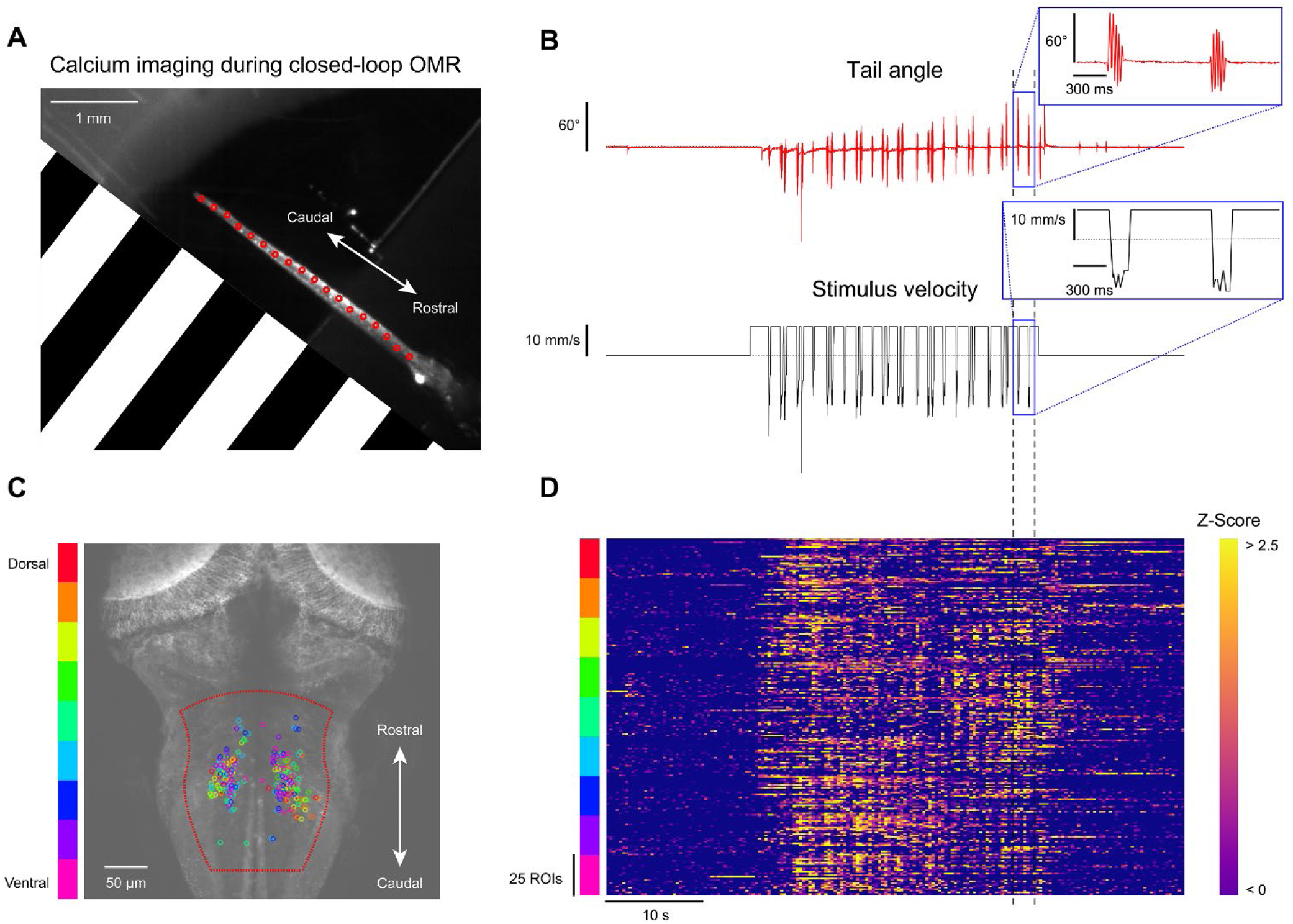
Calcium imaging during closed-loop OMR. One-dimensional closed-loop optomotor gratings were presented to a head-fixed larva while simultaneously performing fast volumetric two-photon calcium imaging. **(A)** Single frame obtained from the behavior camera showing the larva’s tail with the OMR stimulus represented (stimulus spatial frequency was lower than depicted; 450 Hz tracking). The OMR stimulus was presented from the side and travelled from caudal to rostral. **(B)** Tail angle and stimulus velocity across the entire 1-minute trial. Movement of the closed-loop OMR stimulus began at 15 seconds and continued for 30 seconds. **(C)** Maximum intensity z-projection overlaid with neuronal ROIs found in the medial hindbrain. 25 ROIs were selected for each z-plane and color-coded by depth. Automated ROI extraction was performed throughout the region enclosed in dotted red line which helped minimize the detection of non-somatic ROIs. **(D)** Z-scored ROI responses across the trial.

## Discussion

Behavioral tracking systems for larval zebrafish have previously been developed using both commercial and open-source programming languages. ZebraZoom, written in MATLAB, provides methods for high-speed tracking and behavioral classification^16^. Additionally, the open-source Python program Stytra performs high-speed open-loop and closed-loop tracking of larval zebrafish^17^. BonZeb provides a unique tracking solution for zebrafish in that it inherits Bonsai’s framework, uses a high-performance compiled programming language (C#), and operates through a visual programming interface. BonZeb also provides a suite of fully developed visuomotor assays that cover many existing needs for zebrafish researchers.

We demonstrate that BonZeb can perform high-speed online tracking during virtual open-loop predator avoidance, OMR, and prey capture assays. Larvae produced characteristic J-turns, slow approach swims, and eye convergence in response to a small prey-like stimulus in virtual open-loop, consistent with previous findings from naturalistic and virtual prey capture studies^18–20^. In response to optomotor gratings, larvae continually produced turns in the direction of the stimulus, similar to what has been described previously^5,9^. In our predator avoidance assay, larvae displayed rapid directional escape behavior when looming stimuli were presented to either side of the fish in agreement with prior results^8,21^. In the multi-animal OMR assay, we used group behavior to control the direction of optic flow while performing simultaneous online tail tracking for each fish. In the multi-animal prey capture experiment, numerous virtual prey stimuli, presented from below, elicited J-turns toward prey targets across all individuals. As expected, fish displayed much tighter swimming trajectories in the prey assay compared to the long swimming trajectories fish produced in the OMR assay.

The head-fixed assays we developed for BonZeb allow for closed-loop optomotor stimulation in one or two dimensions. In our one-dimensional closed-loop assay, we found that fish produced more bouts more frequently under low gain conditions, with individual bouts having longer durations and higher tail beat frequency compared to bouts generated in high gain conditions. These results agree with previous research that also investigated the effect of visual reafference mismatching on optomotor swimming^10,14^. Our method for head-fixed two-dimensional closed-loop visual feedback builds on the methods of previous research to provide two-dimensional, closed-loop visual feedback to multiple animals simultaneously^11,22^. When presented with randomized initial starting orientations, larvae consistently responded to the OMR stimulus by swimming in the direction of optic flow. Swim bouts were found to align the stimulus orientation toward the animal’s heading direction, in agreement with previous results^11^. We also found that larvae increased their tail beat frequency as the stimulus orientation neared the heading angle.

Bonsai provides a convenient framework for BonZeb to integrate with external devices. We demonstrate this feature with a novel optogenetics assay as well as closed-loop OMR stimulation during calcium imaging. BonZeb’s flexibility in creating tracking pipelines and support of hardware integration allows new users to rapidly develop and implement complex behavioral assays. An exciting future direction is the incorporation of BonZeb’s video acquisition, behavioral tracking, and stimulation capabilities with more complicated 3D visual rendering pipelines using the recently developed BonVision package. This combination would allow the creation of complex immersive environments for virtual open and closed-loop approaches^23^. In summary, BonZeb provides users with a suite of fast, adaptable and intuitive software packages for high-resolution larval zebrafish tracking and provides a diverse set of visuomotor assays to investigate the neural basis of zebrafish behavior.

## Methods

### BonZeb Installation and Setup

A detailed manual for installation, setup and programming in BonZeb can be found on GitHub (https://github.com/ncguilbeault/BonZeb). Briefly, users will need to download and install Bonsai (https://bonsai-rx.org/docs/installation) and the packages required to run BonZeb workflows. Bonsai Video and Bonsai Vision packages are used for video acquisition and video analysis, Bonsai Shaders is used for OpenGL graphics rendering, and Bonsai Arduino is used for communication with Arduino microcontrollers. Packages for BonZeb can be downloaded from Bonsai’s built-in package manager.

### Video Acquisition

BonZeb can be used with a variety of cameras already supported by Bonsai. The FlyCapture package integrates FLIR cameras and the PylonCapture package integrates Basler cameras. As well, the Bonsai Vision package offers access to DirectShow driver-based cameras, such as USB webcams. We have developed packages to incorporate Allied Vision Technologies (AVT) USB 3.0 cameras, Teledyne DALSA GigE cameras, and CameraLink cameras that use Euresys frame grabber boards. Bonsai uses OpenCV for video acquisition and video processing. Thus, our packages use the same implementation of OpenCV structures, formats, and data types. These packages are written in the native C#/.NET language underlying Bonsai’s framework. Our Allied Vision, Teledyne DALSA, and Euresys packages utilize the software development kits (SDK) and .NET libraries (DLLs) provided by the manufacturers to acquire, process, and display frames in Bonsai. Each of our video capture packages offers unique properties for controlling the connected camera’s features. For example, the `VimbaCapture` module from the Allied Vision package allows users to control features of an AVT camera such as frame rate, exposure, black level, gain, and gamma. The output is a `VimbaDataFrame`, containing the image, a timestamp, and a frame ID associated with each image. Similarly, the `SaperaCapture` module performs the same control and output functions for Teledyne DALSA GigE cameras. The properties of each module can be changed dynamically. The ability to modify the camera’s specific features on the fly enables users to quickly optimize the camera’s acquisition properties with respect to their specific imaging requirements. The `MultiCamFrameGrabber` module from the Euresys Frame Grabber package allows users to specify properties such as the `ConnectionType`, `BoardTopology`, and `CameraFile`. While this module does not support dynamic changes to the specific camera’s acquisition features, users can modify the camera file for their camera prior to running Bonsai to adjust the camera’s acquisition settings. The `MultiCamFrameGrabber` module should be compatible with any CameraLink camera that is supported by a Euresys frame grabber board.

### Behavioral Tracking

BonZeb’s package for behavioral tracking is written in C#/.NET and utilizes OpenCV data types for efficient image processing and analysis. We built a `CalculateBackground` module that computes the background as the darkest or lightest image over time (Algorithm 1). This operation works by comparing each individual pixel value of the gray scale input image with the pixel values of a background image contained in memory. The module updates the internal background image only if the pixel value of the input image is greater than the corresponding pixel value of the background image. In this way, if the subject for tracking is darker than the background and the subject has moved, the output of the `CalculateBackground` node will consist of a background image that has successfully removed the subject. By changing the `PixelSearch` property, users can set the module to maintain either the darkest or lightest pixel values in the input image. The optional `NoiseThreshold` parameter can be set to modify the background calculation such that the pixel value of the input image must be greater than or less than the pixel value of the background and some additional noise value. This background calculation method offers advantages over Bonsai’s pre-existing methods for background calculation because it enables rapid background subtraction and only requires a small amount of movement from the animal to obtain a reliable background image.

#### Algorithm 1 Calculation of continuously updated video background

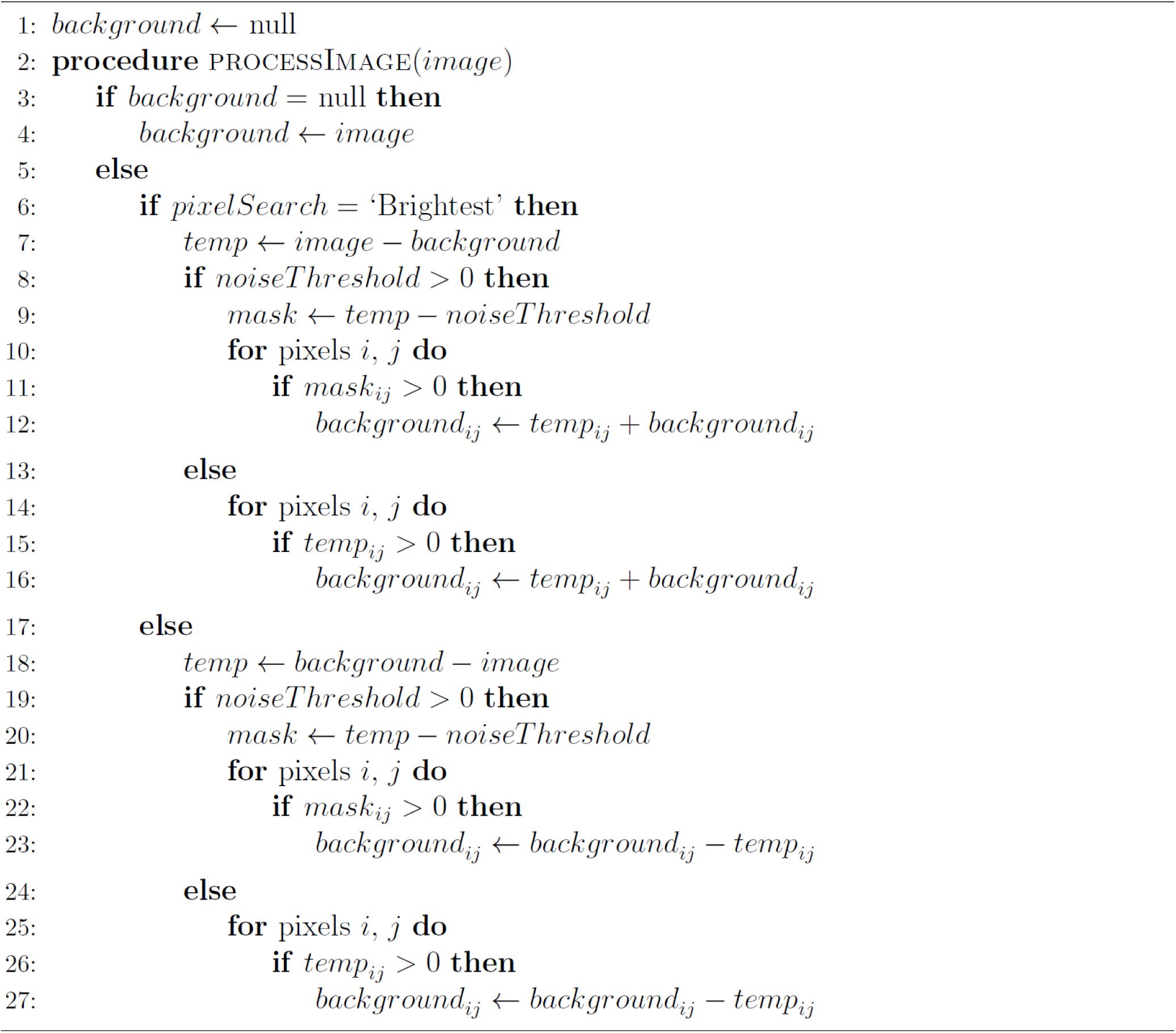

We also provide a module for performing efficient centroid calculation using the `CalculateCentroid` module. This module takes an image as input (usually represented as the background subtracted image) and finds an animal’s centroid using the raw image moments or by finding the largest binary region following a binary region analysis. The method for centroid calculation is set by the `CentroidTrackingMethod` property. The `ThresholdValue` and `ThresholdType` properties set the value of the pixel threshold and the type of threshold applied (binary threshold or inverted binary threshold). If `RawImageMoments` is used, `CalculateCentroid` calculates the centroid using the center of mass of the entire binary thresholded image. The `LargestBinaryRegion` method performs a contour analysis on the thresholded image to find the contour with the largest overall area. This method utilizes the `MinArea` and `MaxArea` properties to discard contours whose area lies outside of the defined range. The result of the `CalculateCentroid` operation using the largest binary region method is the centroid of the contour whose area is the largest within the allowed range of possible areas. In most experiments, the `LargestBinaryRegion` method will be more robust against image noise than the `RawImageMoments` method.

The centroid and the image can then be combined and passed onto the `CalculateTailPoints` module, which fits points along the tail using an iterative point-to-point contrast detection algorithm seeded by the centroid. Two pre-defined arrays are calculated containing the points along the circumference of a circle with a given radius centered around the origin. The first array consists of points along a circle whose radius is equal to the `DistTailBase` property. When the `HeadingDirection` property is negative, the most rostral tail point is found by searching the first array, whose points have been shifted such that the origin is at the centroid. If the `HeadingDirection` is positive, the first tail point is taken as the point in the array whose angle from the centroid corresponds to the angle in the opposite direction provided by the `HeadingDirection`. For subsequent tail points, a second array that contains points along the circumference of a circle with a radius equal to the `DistTailPoints` property is used. A subset of points in this array are calculated and searched for subsequent tail points. This subset of points corresponds to an arc with length determined by the `RangeTailPointAngles` property. The midpoint of the arch is determined by the point in the array where the angle is closest to the angle calculated between the previous two tail points.

Three different contrast-based point detection algorithms can be selected for calculating tail points in the `CalculateTailPoints` module specified by the `TailPointCalculationMethod` property. The `PixelSearch` option is the simplest of the three algorithms and works by taking the darkest or lightest pixel within the array of points. The `WeightedMedian` method involves taking the median of the points whose pixel values are weighted by the difference from the darkest or lightest pixel value. The `CenterOfMass` method calculates the center of mass using the difference of each pixel value from the darkest or lightest pixel value and taking the point in the array that is closest to the center of mass. All three algorithms can calculate the corresponding tail points of the fish efficiently, however, differences between each algorithm will make one or another more desirable depending on the specific needs of the application. While the `PixelSearch` method is faster than the other two methods, the calculated tail points tend to fluctuate more between successive frames. The `WeightedMedian` and `CenterOfMass` methods tend to reduce the amount of frame-to-frame fluctuations but take longer to compute. Despite the minor differences in computational speed, all three algorithms can reliably process images with 1 MP resolution acquired at 332 Hz. Changing the `PixelSearchMethod` property of the `CalculateTailPoints` function will determine whether the algorithm searches for the lightest or darkest pixel values in the image. How many tail points are included in the output depends on the value given to the `NumTailSegments` property, such that the number of points in the output array is the `NumTailSegments` value plus 2. The array is ordered along the rostral-caudal axis, from the centroid to the tip of the tail. Lastly, the `CalculateTailPoints` function contains the optional `OffsetX` and `OffsetY` properties which allows users to offset the final x and y coordinates of the calculated tail points.

The output of the `CalculateTailPoints` is used to extract information about the tail curvature using the `CalculateTailCurvature` module. This function converts an array of points along the tail, *P* = [(*x*_0_, *y*_0_), …, (*x_n_*, *y_n_*)] (ordered from the most rostral to the most caudal), into an array of values Θ = [*θ*_1_, …, *θ_n_*]. The values in the output array correspond to the angles between successive points in the input array normalized to the heading direction. The heading direction (*θ*_h_) is given as the angle between the second point in the array (the most rostral tail point) and the first point (the centroid):

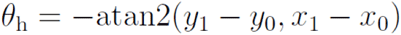

where (*x_i_*, *y_i_*) is the *i*th point along the tail and atan2 is an inverse tangent function that constrains *θ*_h_ such that −π < *θ*_h_ ≤ π. Each point in the array is translated by an amount that is equivalent to translating the centroid to the origin. The heading angle is used to rotate each point in the translated array such that the first angle in the output array is always 0:

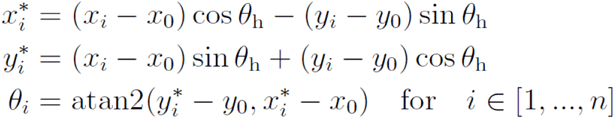

The length of the output array is equivalent to the length of the input array (number of tail points) minus 1. One final computation is performed to normalize the angles. This involves iterating through the entire array and offsetting an angle in the array by 2π if the absolute difference between the current angle and the previous angle is greater than π. This ensures that the angles across all tail segments are continuous and removes disparities between successive angles when relative angles change from −π to π.

The output of the `CalculateTailCurvature` module can then be passed onto the `DetectTailBeatKinematics` module which outputs a `TailKinematics` data structure containing peak amplitude, tail beat frequency, and bout instance. The algorithm used by the `DetectTailBeatKinematics` module (Algorithm 2) works by comparing the current tail angle to an internal memory of the minimum and maximum tail angle over a running window. The difference between the current tail angle and either the maximum or minimum tail angle is used to compare the current tail angle to a threshold, specified by the `BoutThreshold` property. If the difference exceeds this threshold, then the algorithm sets the bout detected value to true and begins searching for inflection points in the stream of input data. The time window, specified by the `FrameWindow` property, determines how many tail angle values to maintain in memory and how many times the counter variable, initialized at the start of a bout and incremented by 1 for each frame a bout is detected, should increment while continuing to search for successive points in the input stream. When a bout is detected, if the difference between the current tail curvature and the maximum or minimum value is less than or greater than the threshold value set by the `PeakThreshold`, then the algorithm begins searching for inflection points in the opposite direction and the peak amplitude is set to the maximum or minimum value. If more than one inflection point has been detected within a single bout, then the tail beat frequency is calculated by dividing the `FrameRate` property by the number of frames between successive peaks in the input data. When the counter variable exceeds the specified `FrameWindow`, the counter variable resets, the bout instance is set to false, and the peak amplitudes and tail beat frequency are set to 0.

#### Algorithm 2 Calculation of tail beat kinematics

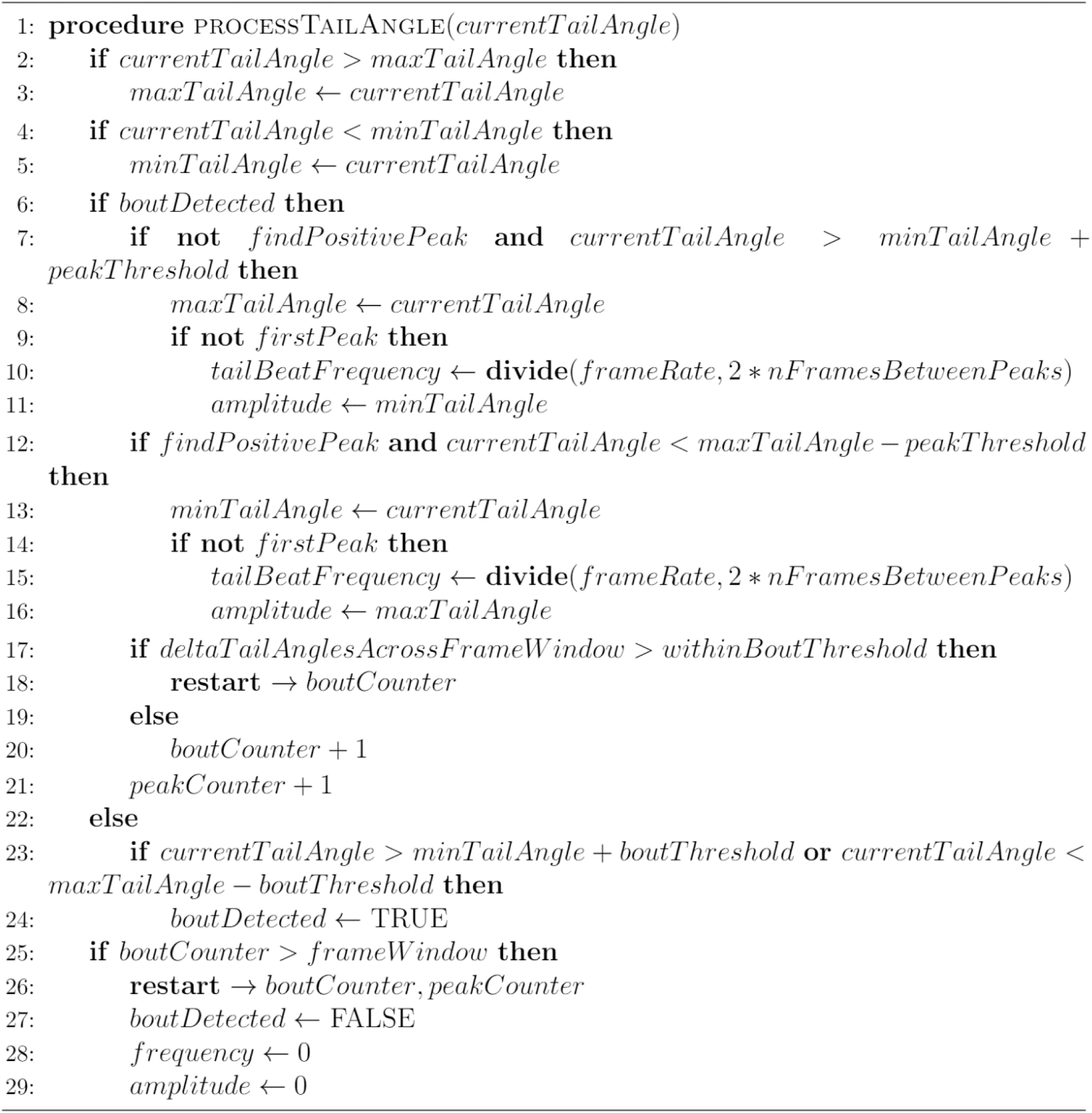

The output of the `CalculateTailPoints` function can be used to find the coordinates of the eyes using the `FindEyeContours` module. The `FindEyeContours` module takes a combination of the array of tail points and a binary image. It processes the image using the output of the `CalculateTailPoints` function to calculate the binary regions corresponding to the eyes. To maintain consistency across our package and the Bonsai Vision package, the output of the `FindEyeContours` module is a `ConnectedComponentCollection`, the same output type as the `BinaryRegionAnalysis` module, which consists of a collection of non-contacting binary regions called “connected components”. The algorithm for finding the eyes starts by performing a contour analysis on the input image, the parameters of which can be specified using the `Mode` and `Method` properties to optimize the contour retrieval and contour approximation methods, respectively. Once all of the contours in the image are acquired, contours whose area lie outside the range of areas provided by the `MinArea` and `MaxArea` properties and whose distance from the centroid of the tracking points lies outside the range of distances given by the `MinDistance` and `MaxDistance` properties, are discarded from further analysis. The remaining contours are then ordered by centering their position around the centroid of the tail tracking points, rotating each contour to face forward with respect to the heading angle, and calculating the absolute difference between the heading angle and the angle between the centroid and the contours’ centroid. The array of contours is ordered in ascending order with respect to the difference between the angle to the centroid and the heading angle. The algorithm continues to discard regions that do not correspond to the eyes by discarding regions that lie outside the `AngleRangeForEyeSearch` property, which is centered around the heading angle. The contours are then ordered by their angles once more, such that the final `ConnectedComponentCollection` consists of the left eye and right eye, respectively. The algorithm also provides the additional `FitEllipsesToEyes` property to allow users to fit ellipses to the eyes.

The output of the `FindEyeContours` module can be combined with the output of the `CalculateTailPoints` function and provided as input to the `CalculateEyeAngles` function to calculate the convergence angles of the left and right eye with respect to the heading angle. For each eye, the function calculates the point *p* = (*x*, *y*) whose distance from the origin is equal to the length of the binary region’s major axis, *l_M_*, and whose angle from the origin corresponds to the orientation of the major axis, *θ*_M_:

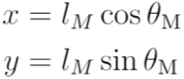

The point is then rotated with respect to the heading angle and the new angle between the point and the origin, *θ*^∗^, is calculated:

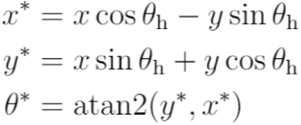

The orientation of the binary region’s major axis orientation is always between 0 and π. This poses a problem for calculating the orientation of the eyes with respect to the heading direction, which is between 0 and 2π, because the orientation of the eyes needs to be in a consistent direction with respect to the heading angle. The `CalculateEyeAngles` module accounts for this by checking which direction the angle of the eye faces with respect to the heading direction and then offsets the angle by π if the eye angle is initially calculated in the opposite direction. Alternatively, the heading angle, derived from the `CalculateHeadingAngle` module, can be substituted for the array of tail points as input into the `CalculateEyeAngles` function to determine the angles of the eyes with respect to heading angle if only the centroid and heading angle data are available. A combination of the array of tail points, generated by the `CalculateTailPoints` operation, and the eye contours, generated by the `FindEyeContours` function, can be passed as input to the `CalculateHeadingAngle` module to calculate a more accurate representation of the heading angle using the angle between the centroid and the midpoint between the centroids of the two eyes. The `CalculateHeadingAngle` module, which tracks the cumulative heading angle over time, also has an option to initialize the heading angle to 0 at the start of recording, which can be set with the `InitializeHeadingAngleToZero` property.

### Visual Stimulus Library

We used the Bonsai Shader package and OpenGL to compile a library of visual stimuli that can be used for closed-loop and virtual open-loop assays. Virtual open-loop visual stimuli are updated based on the position and heading direction of the animal. The library contains the following stimuli: looming dot, optomotor gratings, phototaxic stimuli, optokinetic gratings, single or multiple prey-like small dots, and whole field luminance changes (black and white flashes). Additionally, we have developed a method for closed-loop stimulation of head-restrained animals using methods we have developed for analyzing the tail curvature of head-restrained larval zebrafish to estimate changes in position and heading direction. We demonstrate how to use this method to provide head-fixed larval zebrafish with one-dimensional or two-dimensional closed-loop OMR stimulation. Our method for closed-loop OMR stimulation in head-fixed animals can be easily extended to work with other stimuli such as the looming dot stimulus, prey stimulus, and multi prey stimulus.

Calibrating the display device to the camera involves identifying the areas of the display that correspond to the camera’s field of view (FOV). To do so, users need to align the edges of a projected box onto the edges of the camera’s FOV. Users can either remove the IR filter on the camera to allow the image from the display to show up on the camera, or use an object, such as a translucent ruler, as a common reference to align the edges of the projected box to the edges of the camera’s FOV. We developed a fast and reliable method to map this area using the `DrawRectangle` module, which allows users to draw a rectangle onto an input image and outputs a `CalibrationParameters` construct. The outputs of the `CalibrationParameters` can then be mapped directly onto a stimulus to modify the area the stimulus is presented to. By drawing a simple rectangle shader onto a specific area of the display, the user can then determine the area of the projector that corresponds to the camera’s FOV.

### Virtual Open-Loop and Closed-Loop Setup

We implemented an established optomechanical configuration to record behavior of fish and project computer generated stimuli from below^5,8,9^. This setup included a projector for visual stimulation, an infrared backlight for fish illumination, a cold mirror and a high-speed camera equipped with a long-pass filter. A Sentech STC-CMB200PCL CameraLink camera (Aegis Electronic Group, Inc.) was used. The camera was integrated using a GrabLink Full XR Frame Grabber (Euresys Inc.). This configuration allowed us to acquire 1088 x 1088 resolution frames at 332 Hz. We also used the same configuration to acquire 640 x 480 resolution frames at 700 Hz. We used a fixed focal length (30 mm) lens (Schneider-Kreuznach) equipped with a long-pass filter (Edmund Optics) to block out the visible light from back-projected stimuli presented from below the camera. A high-definition, 1920 x 1080, 60 Hz, DLP pico projector (P7, AAXA Technologies Inc.) was used for visual stimulation and a 25 x 20 cm cold mirror (Knight Optical) reflected visible light from the projector upwards to the bottom of the platform. An 850 nm LED array (Smart Vision Lights) provided tracking light and was placed underneath the cold mirror. The platform consisted of a 6 mm thick transparent polycarbonate sheet (20 cm x 22 cm), with mylar diffusion paper placed on top for back projection. The workstation computer was equipped with an Intel Core i9-9900K processor, an ASUS ROG GeForce GTX 1060 graphics card, and 64 GB of RAM.

### Virtual Open-Loop Looming Stimulation

Fish were placed in a 6 cm watch glass inside a 10 cm petri dish. Both the petri dish and watch glass were filled with filtered system. The water was 5 mm in depth. Looming stimulation trials were 20 seconds long. The angular size to speed ratio (l/v) of the looming stimulus was 90 ms. At the start of the trial, the looming stimulus was shown for 10 seconds without expansion. After 10 seconds, the looming stimulus began to expand after which the size was truncated at 120° ∼13 seconds into the trial. Looming stimuli were positioned 90° to the left or right of the fish and 1 cm away from the centroid. The position of the looming dot was calculated using the position and heading angle of the fish. Escapes were automatically detected by selecting the bouts whose max velocity exceeded 15 cm/s. Individual escape trajectories were calculated by taking the change in coordinates from the start of the bout. The coordinates were rotated so that the heading angle at the start of the bout faced the same direction in all cases. We calculated the change in heading from the start of the bout to the time the first change in peak amplitude was detected to obtain the max initial heading angle.

### Virtual Open-Loop OMR Stimulation

We used sinusoidal gratings with a spatial frequency of 1 cm and a velocity of 1 cm/s. The orientation of the optomotor gratings was calculated using the heading angle of fish. The position of the fish was used to calculate the pivot point for changing the orientation of the gratings. Trials were 1-minute long and consisted of continuous OMR stimulation 90° leftward or rightward.

### Virtual Open-Loop Prey Stimulation

Virtual prey stimuli were presented in 1-minute trials. Prey stimuli moved along an arc of 120° spanning 60° to the left and right of the fish in sinusoidal motion. The prey stimuli were positioned 5 mm away from the fish and were < 1 mm in diameter. The position of the prey was calculated using the centroid coordinates and the heading angle.

### Behavioral Clustering of Virtual Prey Responses

We used hierarchical clustering to separate virtual prey responses into different bout classes. We calculated four kinematic parameters for each bout (mean tail beat frequency, bout integral, bout standard deviation, and max tail amplitude). For the clustering analysis, we ignored the specific left-right direction of the bout by taking the absolute value of the bout integral and max tail amplitude. We calculated the average silhouette score across bouts using a range of models with clusters from 2 – 10. We used the model which produced the maximum silhouette score. When calculating the mean tail angle for each bout, bouts were normalized and plotted such that the bout integral was positive. The identities of the bout clusters we resolved (forward swims, routine turns, and J-turns) were determined by comparing the results of our analysis to the results of previous literature^24,25^. We then decomposed the normalized bouts in the J-turn cluster into unnormalized left-biased and right-biased J-turns using the sign of the bout integral to determine the direction. We calculated the prey yaw angle by taking the position of the virtual prey stimulus with respect to the position and heading angle of the fish at the start of each J-turn.

### Multi-Animal Tracking

For multi-animal tracking with optomotor stimulation, we used sinusoidal gratings with a spatial frequency of 1 cm and a velocity of 1 cm/s. We built a simple protocol that calculated the center of mass of the group by taking the average of all tracked centroids. When the group’s center of mass entered the leftmost quarter of the arena, the direction of the OMR stimulus updated to move rightwards. When the center of mass was detected in the rightmost quarter of the arena, the direction of the stimulus changed to travel leftwards.

For multi-animal tracking with virtual prey, we projected 6 virtual prey stimuli ∼1 mm in diameter. Each virtual prey was programmed to move in a paramecia-like manner, such that prey would move in a specific direction for a randomly determined duration, followed by a brief pause and then movement in a different direction. Trials were 1-minute long. J-turns in response to virtual prey stimulation were manually identified in the video by a human observer.

To calculate the identities of the fish, we developed a custom post-processing analysis pipeline that could determine the identities in retrospect. We computed the Euclidean distance between all centroids at successive points in time and performed a linear sum assignment to minimize the total distance between all pairs of centroids. When fish came into contact, their centroids merged into a single centroid during online calculation. This led to fewer centroids than the expected number of centroids calculated at those time points. Under these conditions, we calculated which centroid at the current time point minimized the distance between the unassigned centroid in the previous time point and all current centroids. We then assigned this centroid to the identity of the unassigned centroid.

### One-Dimensional Closed-Loop OMR

The one-dimensional closed-loop OMR assay we implemented was based on a previous study^10^. The one-dimensional closed-loop sensory feedback was calculated using the following transformation,

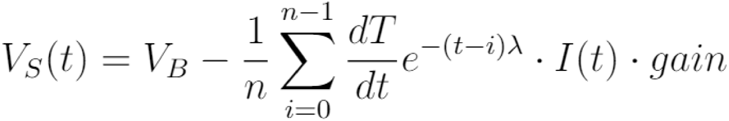

where the stimulus velocity (*V_s_*) at time *t* was the result of the baseline velocity (*V_B_*) minus the swim vigor. The swim vigor was calculated as the derivative of the tail angle (*dT/dt*) integrated over a 15 ms time window with a decay constant of 9 ms. The exponential function served to simulate the effects of acceleration and deceleration during bursts and glides. The tail angle derivative was multiplied by the instance of a detected bout (*I*) at time *t*, where *I*(*t*) equals 1 when a bout is detected and 0 otherwise. Finally, the swim vigor was multiplied by a constant *gain* factor.

For this head-fixed preparation, fish were mounted in 2% low-melting point agarose inside of a 4 cm petri dish. Fish were positioned roughly 5 mm above the bottom of the dish. After letting the agarose set for ∼5-10 minutes, we added filtered system water to a depth of 8 mm. We freed the tails of the fish by cutting away the agarose caudal to the swim bladder with a razor blade. Fish recovered from mounting for 4 – 12 hours in the agarose before testing.

Fish were stimulated with a square wave grating with a spatial frequency of 10 mm. We initially presented fish with a static grating for 30 seconds. Following this initial waiting period, a 10 second long trial was initiated where the gratings moved at a baseline velocity of 10 mm/s and the fish’s swim vigor was used to simulate closed-loop visual feedback. After the 10 second trial, the motion of the gratings ceased and a 30 second no motion inter-trial-interval commenced. For each trial, the gain factor was set to either 0.5 (low), 1 (medium), and 1.5 (high) to provide varying degrees of perceived visual feedback. The calculation of swim vigor was set such that in trials with a medium gain factor of 1, a tail beat frequency of 20 Hzapproximated a 20 mm/s forward translation, resulting in a final −10 mm/s stimulus velocity. Under low gain conditions (0.5), a tail beat frequency of 20 Hz only resulted in 10 mm/s forward translation, transforming the final stimulus velocity into 0 mm/s. When the gain was high (1.5), a bout with a 20 Hz tail beat frequency resulted in 30 mm/s forward motion, generating a −20 mm/s visual feedback. Each gain value was presented in three consecutive trials, before switching to a different gain. The first three trials had a medium gain, followed by low gain and then high gain. After the first nine trials, fish were presented with medium gain, followed by high gain and then low gain. Blocks of 18 trials were repeated three times for a total of 54 trials for each fish.

Similar to previous work^10^, we used normalized values to determine the relative change to a fish’s behavior under different gain conditions. To do this, the bout kinematics for each fish were divided by the average value for that fish across all trials. We then averaged the normalized kinematics of a single fish across all trials with the same gain value.

### Multi-Animal Two-Dimensional Closed-Loop OMR

The design of the two-dimensional closed-loop OMR assay was inspired by work in a previous study^11^. We used the following transformation to update the velocity of the stimulus:

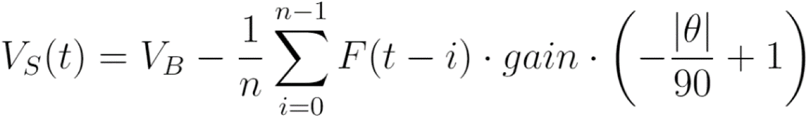

The stimulus velocity (*V_s_*) at time *t* was calculated as the base velocity (*V_B_*) minus the tail beat frequency (*F*) multiplied by a constant *gain* factor. The velocity of the OMR stimulus along a single dimension was directly proportional to the online calculated tail beat frequency. A moving average (box car filter) was applied to the tail beat frequency. This was used to simulate the effects of bursting and gliding during swimming. The velocity of the stimulus also depended on the orientation of the stimulus (*θ*) relative to the heading direction, where *θ* ∈ [-180°, 180°). This term reaches a maximum value of 1 when the stimulus is perfectly aligned with the heading angle at 0°. We calculated the orientation of the stimulus using the following equation,

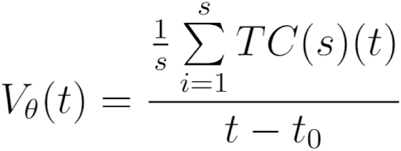

The angular velocity of the OMR stimulus (*V_θ_*) was a function of the mean tail curvature (*TC*) along all calculated tail segments (*s*) at a given time *t* weighted by the inverse of time since the onset of the swimming bout (*t*_0_). The equation for angular velocity weighs the amplitude of the mean tail curvature with the relative length of the swimming bout, such that the initial amplitudes of the swimming bout produced larger changes to the angular velocity compared to subsequent amplitudes.

On a 10 cm petri dish, we mounted one fish at a time (4 total) in a 2 x 2 grid with all fish facing the same direction. Once the agarose was set for all fish, filtered system water was added. We inserted a laser-cut IR-transmitting black acrylic piece shaped like a cross to separate fish into quadrants. This was to ensure that fish could not see neighboring stimuli. We tested the four fish simultaneously, each presented with its own two-dimensional closed-loop OMR stimulus. We used a sinusoidal grating with a spatial frequency of 10 mm. We used a gain value of 1 for all trials. In a single test session, larvae were randomly presented with one of 8 grating orientations that were equally positioned around the unit circle [-180° to 135°]. After an initial 1-minute period with static gratings, the gratings moved at a velocity of 10 mm/s for 30 seconds, after which both stimulus velocity and stimulus orientation were updated based on the fish’s behavior. Each initial starting orientation was presented to the fish four times for a total of 32 trials.

For calculating the probability density of stimulus orientations over time, we included all of the trials across all fish. The distribution of values for each time bin were determined by taking a snapshot of values at the point in time when the time bin started. The distribution of stimulus orientations at each point in time was normalized across trials. Stimulus orientations were binned into 45° samples. To compare the effects of stimulus orientation on bout kinematics, we used the stimulus orientation at the start of each bout. We grouped bouts based on their stimulus orientation into bins of 45° and performed a one-way ANOVA on the binned bouts. Finally, we used a polynomial regression to model the relationship between kinematics and stimulus orientation. A range of models were tested with degrees ranging from 1 – 10, and the model with the greatest change in explained variance was taken as the best model.

### Optogenetic Assay

The optogenetic stimulation setup used a custom LED array arranged in a 3 x 3 grid of high-powered blue LEDs (470 nm) that produced 0.9 mW/mm^2^ at the stage. The LED array was placed above and to the side of the platform and faced a 20 cm x 20 cm cold mirror (Knight Optical) angled at a 45° angle to direct the short wavelength light coming from the blue LED array towards the platform while transmitting IR light to the camera. A Basler ace acA1920-155uc (Basler AG) camera equipped with a lens (Edmund Optics) and long-pass filter (Edmund Optics) was used to record behavior at 200 Hz with a resolution of 1200 x 750. Four IR LED panels, each consisting of 12 LEDs (850 nm) provided behavior illumination (Autens IR Illuminator, Amazon). An Elegoo UNO R3 board connected via USB to the workstation was used to control the timing of LED illumination (50 Hz; 10 ms pulse width). The workstation was equipped with an Intel Core i7-6700K CPU, an EVGA GeForce GTX 1060 GPU, and 32GB of RAM. The platform of the behavioral assay consisted of a 19.5 x 14.5 cm transparent polycarbonate sheet with mylar diffusion paper placed on top. We tested 10 fish simultaneously by allowing them to freely swim in individual wells of 2.5 x 1.5 cm with a depth of 5 mm using a custom multi-well behavioral arena. All fish were tracked simultaneously in a multi-well plate, while providing timed stimulation light with a high-powered 470 nm LED array modulated by an Arduino microcontroller. We synchronized the output of the Arduino to the camera’s frame rate (200 Hz) and delivered pulsed optical stimulation (50 Hz, 10 ms pulse width). We used fish expressing channelrhodosin-2 in glutamatergic neurons: *Tg(vglut2a:Gal4)^uot13^; Tg(UAS:ChR2(H134R)-mCherry)^s1986t^*. For each experiment, 5 expressing and 5 non-expressing clutch-mates were placed pseudo randomly into one of the wells of the multi-well arena.

Each experiment was 20 minutes long with stimulation given in periods of 1 minute with 1-minute intervals of no stimulation. The beginning and end of the experiment had 30 second periods without stimulation. To calculate the mean velocity over stimulation periods, we averaged across all 2-minute periods for each fish and then averaged across fish to obtain the mean velocity over time for each group. We then calculated the cumulative distance travelled for each fish across all periods of stimulation ON and stimulation OFF separately. We used a mixed ANOVA design to compare experimental versus control animals (between group factor) and distance travelled during stimulation ON versus stimulation OFF (within group factor). We then conducted pairwise t-tests to compare the effects of stimulation ON/OFF within each group.

### Closed-Loop OMR During Calcium Imaging

We modified the head-fixed closed-loop behavioral setup described above so that we could implement the same techniques under a two-photon microscope. We illuminated the tail of the fish using a high-powered 850 nm LED (Thorlabs), placed above the stage and angled at 45°. We placed a small, 5 x 5 cm gold protected mirror, underneath the fish and angled it at 45° to reflect light towards the behavior camera. We used a Genie Nano M640 NIR camera (Teledyne DALSA), zoom lens (Navitar) and an IR long-pass filter (ThorLabs) to acquire video at 450 Hz with a resolution of 640 x 480. Visual stimuli were presented using a 1920 x 720, 60 Hz, laser pico projector (MP-CL1A, Sony) onto a 10 x 10 cm screen made by adhering mylar diffusive paper to a 3D printed rectangular frame. A TTL signal from the microscope upon the start of scanning was sent to an Arduino Uno to synchronize imaging, visual stimulation, and the behavior camera. The Arduino, projector, and camera were controlled by a workstation with an Intel Core i7-4790 CPU, a Radeon R7 250 GPU, and 32GB of RAM.

Fish expressing a pan-neuronal calcium indicator *Tg*(*elavl3:GCaMP6f*)*^jf1^* were embedded in 2% low melting point agarose with the caudal half of the tail free. To drive optomotor swimming, a square wave grating with a velocity of 1 cm/s and spatial frequency of 1 cm was projected onto a screen mounted parallel to the larval fish. We registered the scan trigger signal of the two-photon microscope with the analog pin of an Arduino. The state of the pin was monitored inside the Bonsai workflow and the rising edge of the analog signal was used to initiate data collection as well as presentation of a static OMR stimulus. After 15 seconds, the OMR stimulus began to move for 30 seconds. The stimulus velocity was updated dynamically using the feedback equation above with a gain of 1.

### Closed-Loop Round-Trip Latency

Round-trip tests were conducted using a head-fixed larval zebrafish with closed-loop OMR stimulation. We used the same protocol described in the one-dimensional closed-loop methods section with the following modification. We rendered a black rectangle on top of the gratings that covered an area slightly larger than the fish’s head and body. In the upper left corner of this rectangle, we projected a square that flashed white when a bout was detected and black otherwise. The same digital signal from the bout detection algorithm was sent simultaneously to an Arduino-controlled LED that was visible to the camera to turn ON when a bout was detected. We used the same closed-loop feedback calculation as for the one-dimensional OMR experiment. We calculated the number of frames between when a bout was detected in the image and when the LED turned ON, as well as when a bout was detected and when the square from the projector flashed white. We then converted the number of frames to a time delay in ms.

### Multi-Animal Tracking Performance

For single animal tracking, we used the same virtual open-loop OMR assay described above and for multi-animal tracking with group sizes of 5, 10, 15 & 20 we used the same procedure as the multi-animal OMR assay. We conducted 3 separate tests for each group size. Each test was 2 minutes. Tracking data points were timestamped online. We then calculated the difference in timestamps between successive data points. For percent accuracy, we calculated the percentage of tracking data where no fish-to-fish contacts were detected. We used the same identity calculation method described in the multi-animal tracking section. The percent accuracy was calculated with the following equation,

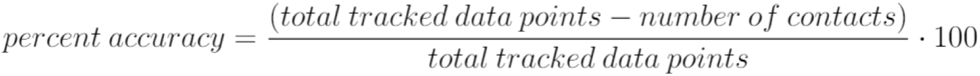

Finally, we calculated the average of the percent accuracy across all tests.

### Quantification and Statistical Analyses

Statistical analyses of the data were performed in Python using custom scripts and imported functions from various python packages. NumPy and Pandas were used for processing data, organizing data, and performing numerical calculations. SciPy, Pingouin, and statsmodels were used for statistical analyses. Scikit-learn was used for clustering. Matplotlib and OpenCV were used in Python for generating data figures and plotting trajectories. Bonsai and BonVision was used to overlay text, tracking points and visual stimuli over raw video. Blender was used for 3D modelling. Raw data collected from BonZeb included tail curvature, heading angle, stimulus angle, position coordinates, bout instances, tail beat frequency, and peak amplitudes. We calculated the start and ends of individual bouts using the rising edge and falling edge of the bout detected signal. 15 ms before and after the start and end of each bout was included for analyses. We calculated the tail angle as the average curvature of the last 3 tail segments. All analyses scripts can be supplied upon request.

### Animals

All experiments were approved by the University of Toronto Local Animal Care Committee and adhered to the governance of the Canadian Council on Animal Care. The study was carried out in accordance with the ARRIVE guidelines. Zebrafish were raised and bred at 28° C on a 14 hr light/10 hr dark cycle. Larvae were fed paramecia starting from 4 days post fertilization (dpf). A zebrafish strain recently obtained from large breeding ponds in Malaysia was used for free-swimming virtual open-loop and head-fixed closed-loop experiments (6-8 dpf). Larval fish (6-8 dpf) expressing channelrhodopsin-2 in glutamatergic neurons were used for the optogenetic assay: *Tg(vglut2a:Gal4)^uot13^; Tg(UAS:ChR2(H134R)-mCherry)^s1986t^*. *Gal4^uot13^* was generated by injecting Cre RNA into one-cell stage embryos from a *vglut2a:loxP-mcherry-loxP-Gal4* line^26^. *Tg(elavl3:GCaMP6f)^jf1^* larvae (6-8 dpf) were used for closed-loop OMR calcium imaging experiments.

## Acknowledgements

This research was supported by an NSERC doctoral scholarship (N.G.), NSERC Discovery grant (T.T.), a Human Frontier Science Program grant (RGY0079/201, T.T.), University of Toronto Connaught New Researcher Award (T.T.) and funding from the Canadian Foundation for Innovation and the Ontario Research Fund (T.T.).

## Author Contributions

N.G. and T.T. devised the project. N.G. and T.T. designed and constructed the closed-loop and virtual open-loop behavioral setup. N.G. wrote the behavioral tracking software. N.G. conducted and analyzed the virtual open-loop, closed-loop, and multi-animal tracking experiments. N.G. and J.G. wrote the video acquisition software. T.T., J.G. and M.M. designed and built the calcium imaging setup. M.M. collected the calcium imaging data. N.G., J.G. and M.M. analyzed the calcium imaging data. T.T., N.G. and I.T. designed and constructed the optogenetics setup. I.T. collected the optogenetics data and N.G. analyzed the optogenetics data. N.G. and T.T. prepared figures and wrote the manuscript.

## Additional Information

Supplementary videos and manual can be found at https://github.com/ncguilbeault/BonZeb. The authors declare no competing financial interests.

## References

1. von Holst, E. & Mittelstaedt, H. The reafference principle. Interaction between the central nervous system and the periphery. In Selected Papers of Erich von Holst: The Behavioural Physiology of Animals and Man (Transl. by Martin, R., ed.) 1. 39–73 (University of Miami Press, 1950).

2. Reichardt, W. & Poggio, T. Visual control of orientation behaviour in the fly: Part I. A quantitative analysis. Q. Rev. Biophys. 9, 311–375 (1976).

3. Weber, T., Thorson, J. & Huber, F. Auditory behavior of the cricket. J. Comp. Physiol. 141, 215–232 (1981).

4. Strauss, R., Schuster, S. & Gotz, K.G. Processing of artificial visual feedback in the walking fruit fly Drosophila melanogaster. J. Exp. Biol. 200, 1281–1296 (1997).

5. Orger, M.B., Kampff, A.R., Severi, K.E., Bollmann, J.H. & Engert, F. Control of visually guided behavior by distinct populations of spinal projection neurons. Nat. Neurosci. 11, 327–33 (2008).

6. Dombeck, D.A. & Reiser, M.B. Real neuroscience in virtual worlds. Curr. Opin. Neurobio. 22, 3–10 (2012).

7. Lopes, G. et al. Bonsai: An event-based framework for processing and controlling data streams. Front. Neuroinform. 9, 7 (2015).

8. Dunn, T.W. et al. Neural circuits underlying visually evoked escapes in larval zebrafish. Neuron 89, 613–628 (2016).

9. Naumann, E.A. et al. From whole-brain data to functional circuit models: The zebrafish optomotor response. Cell 167, 947–960 e20 (2016).

10. Portugues, R. & Engert, F. Adaptive locomotor behavior in larval zebrafish. Front. Syst. Neurosci. 5, 72 (2011).

11. Jouary, A., Haudrechy, M., Candelier, R. & Sumbre, G. A 2D virtual reality system for visual goal-driven navigation in zebrafish larvae. Sci. Rep. 6, 34015 (2016).

12. Dunn, T.W. et al. Brain-wide mapping of neural activity controlling zebrafish exploratory locomotion. eLife 5, e12741 (2016).

13. Giovannucci, A. et al. CaImAn an open source tool for scalable calcium imaging data analysis. eLife 8, e38173 (2019).

14. Ahrens, M.B. et al. Brain-wide neuronal dynamics during motor adaptation in zebrafish. Nature 485, 471–477 (2012).

15. Severi, K.E., Böhm, U.L., & Wyart, C. Investigation of hindbrain activity during active locomotion reveals inhibitory neurons involved in sensorimotor processing. Curr. Biol. 23, 843–849 (2013).

16. Mirat, O., Sternberg, J.R., Severi, K.E. & Wyart, C. ZebraZoom: An automated program for high-throughput behavioral analysis and categorization. Front. Neural Circuits 7, 107 (2013).

17. Stih, V., Petrucco, L., Kist, A.M. & Portugues, R. Stytra: An open-source, integrated system for stimulation, tracking and closed-loop behavioral experiments. PLOS Comput. Biol. 15, e1006699 (2019).

18. Budick, S.A. & O’Malley, D.M. Locomotor repertoire of the larval zebrafish: Swimming, turning and prey capture. J. Exp. Biol. 203, 2565–2579 (2000).

19. Bianco, I.H., Kampff, A.R. & Engert, F. Prey capture behavior evoked by simple visual stimuli in larval zebrafish. Front. Syst. Neurosci. 5, 101 (2011).

20. Mearns, D.S., Donovan, J.C., Fernandes, A.M., Semmelhack, J.L. & Baier, H. Deconstructing hunting behavior reveals a tightly coupled stimulus-response loop. Curr. Biol. 30, 54–69 (2019).

21. Temizer, I., Donovan, J.C., Baier, H. & Semmelhack, J.L. A visual pathway for looming-evoked escape in larval zebrafish. Curr. Biol. 25, 1823–1834 (2015).

22. Chen, X. et al. Brain-wide organization of neuronal activity and convergent sensorimotor transformations in larval zebrafish. Neuron 100, 876–890 e5 (2018).

23. Lopes, G. et al. Bonvision – an open-source software to create and control visual environments. bioRxiv (2020).

24. Marques, J.C., Lackner, S., Felix, R., & Orger, M.B. Structure of the zebrafish locomotor repertoire revealed with unsupervised behavioral clustering. Curr. Biol. 28, 181–195 (2018).

25. Fernandes, A.M. et al. Neural circuitry for stimulus selection in the zebrafish visual system. Neuron (2020).

26. Satou, C. et al. Transgenic tools to characterize neuronal properties of discrete populations of zebrafish neurons. Development 140, 3927–3931 (2013).

