## Supplementary Material for "BonZeb: Open-source, modular software tools for high-resolution zebrafish tracking and analysis"

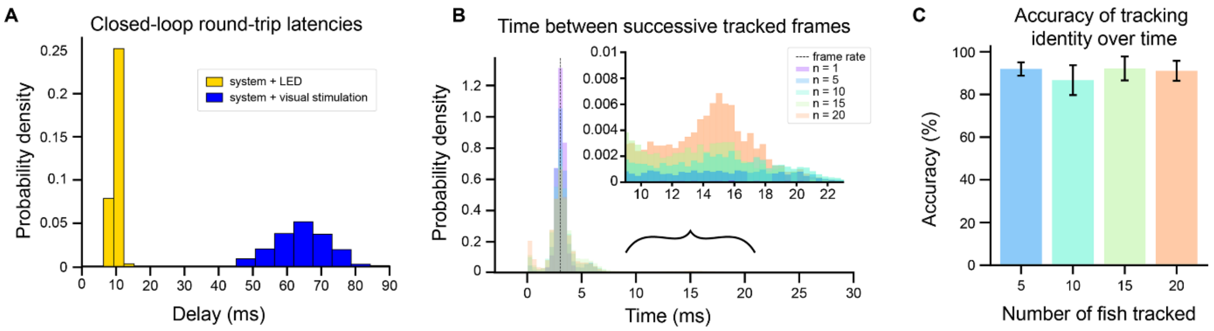


**Supplementary Figure 1. Performance metrics. (A)** Round-trip latencies for a head-fixed fish in a closed-loop OMR environment with a gain of 1. The round-trip was calculated as the difference in time between when a bout was detected and when a corresponding change to an LED (yellow) or visual stimulus (blue) was detected in the video. The probability density of each distribution was normalized by dividing the distribution by the maximum binned value within each distribution. **(B)** Distribution of the time between successive tracked frames with multi-animal tracking for different group sizes. The dashed line represents the expected time for a frame rate of 332 Hz. Buffering of internal resources leads to some variance in the distribution of time data for a single animal when no frames are dropped, with the mean centered around the theoretical frame rate. As the number of tracked fish increases, the data become more skewed towards larger time values. **(C)** Accuracy of tracking identities over time during multi-animal tracking. Accuracy was calculated as the total number of tracked data points minus the number of data points when fish-to-fish contact was detected all divided by the total number of tracked data points.

**Supplementary Video 1. Looming escape.** An example of an escape response generated during virtual open-loop looming stimulation where the looming dot fully expands. Video is shown at 0.2x speed. Raw video was captured at 332 Hz at 1088 x 1088 resolution. Tracking points and looming stimulus were overlaid onto raw video post-processing. Top graph shows tail angle ($^{o}$) over time. Bottom graph shows heading angle ($^{o}$) over time.

**Supplementary Video 2. Looming escape at 700 Hz.** An example of an escape response generated during virtual open-loop looming stimulation where the looming dot has a truncated expansion and follows the fish. Video is shown at 1x speed. Raw video was captured at 700 Hz at 640 x 480 resolution. Tracking points and looming stimulus were overlaid onto raw video post-processing. Top graph shows tail angle ($^{o}$) over time. Bottom graph shows heading angle ($^{o}$) over time.

**Supplementary Video 3. Virtual open-loop OMR.** An example video of a fish producing optomotor responses to drifting black and white sinusoidal gratings in virtual open-loop. Video is shown at 1x speed. Raw video was captured at 332 Hz at 1088 x 1088 resolution. Tracking points and the optomotor stimulus were overlaid onto the raw video post-processing.

**Supplementary Video 4. Virtual prey capture with a moving prey target.** An example of a larval zebrafish engaging in multiple hunting episodes towards a moving virtual prey stimulus. Video is shown at 0.5x speed. Raw video was captured at 332 Hz at 1088 x 1088 resolution. Virtual prey stimulus was overlaid onto raw video post-processing.

**Supplementary Video 5. Virtual prey capture with a locally fixed prey target.** An example of multiple J-turns in rapid succession during virtual open-loop prey stimulation with the virtual prey stimulus fixed to a specific location in the visual field. Video is shown at 0.5x speed. Raw video was captured at 168 Hz at 1280 x 1024 resolution. Raw video captured both the fish and the stimulus. Tracking points were overlaid onto video post-processing.

**Supplementary Video 6. Multi-animal tracking with OMR.** An example of multi-animal tracking with 9 fish during OMR stimulation. OMR stimulus is not shown. Video is shown at 1x speed. Video was captured at 332 Hz at 1088 x 1088 resolution. Tracking points were overlaid onto the raw video in real-time. Fish identities were overlaid onto the video post-processing.

**Supplementary Video 7. Closed-loop OMR with multiple gains.** An example composite video showing the same head-fixed fish stimulated with closed-loop OMR under three different gain values. Each video is shown at 1x speed. Videos were captured at 332 Hz at 1088 x 1088 resolution. Raw video captured both the fish and the stimulus. The stimulus appears low contrast due to the IR backlight. Composite video and text were overlaid post-processing.
